# Mechanochemical synchronization of force oscillations in connective tissue models

**DOI:** 10.64898/2026.10.18.759356

**Authors:** David Böhringer, Leah Vogler, Christoph Mark, Pamela L. Strissel, Jonas Röder, Delf Kah, Lovis Schween, Olivier Zajac, Annamaria Härtl, Lena Fischer, Silvia Budday, Richard Gerum, Harald Herrmann, Lilli Winter, Gerhard Wiche, Wolfgang H. Goldmann, Giovanni Cappello, Danijela Matic Vignjevic, Martial Balland, Aldo R. Boccaccini, Reiner Strick, Ingo Thievessen, Thomas Boudou, Ben Fabry

## Abstract

Cells in a tissue coordinate their behavior collectively by mechanical and chemical feedback with their surroundings. To study how such feedback mechanisms interact, we embed multicellular spheroids from mouse and primary human fibroblasts of different origin in 3D collagen matrices. Immediately after embedding, spheroids apply steadily increasing, radially symmetric, inward-directed forces to the surrounding matrix. Typically, within a few hours, collective force oscillations emerge with an average time interval of *≈*30 min. Similar force oscillations also arise in engineered, parallel-aligned microtissues attached to flexible pillars. Force oscillations propagate between neighboring cells as a mechanical wave, accompanied by calcium waves that travel at comparable speeds of several hundred µm/min. Blocking gap junctions or mechanosensitive Piezo1 channels impairs these oscillations, indicating that calcium signaling contributes to their propagation. Their amplitude and frequency are regulated by integrin-mediated feedback in actin-rich protrusions and by ERK signaling, which also controls collective invasion into the surrounding matrix. Neighboring spheroids mechanically synchronize their force oscillations in phase and frequency over distances of up to *≈*2 mm without direct contact. Together, our results show that oscillatory mechanochemical feedback enables multicellular systems to coordinate force generation and invasive migration across cellular and millimeter length scales in 3D matrices.

## Introduction

In tissues, cells are mechanically and chemically coupled both through the extracellular matrix and through direct cell-cell contacts. Through this coupling, forces generated by one cell deform surrounding cells, which can convert these mechanical cues into biochemical signals and adjust their own contractility, adhesion, or cytoskeletal turnover. This reciprocal feedback helps coordinate collective migration, tissue organization and remodeling, e.g., during wound healing, morphogenesis, or cancer progression (1–6). One well-studied example of such mechanochemical feedback is the propagation of extracellular signal-regulated kinase (ERK) activity waves. In epithelial monolayers, local cell contractions stretch neighboring cells and activate ERK.

ERK then induces a delayed contractile response, which stretches the next cells and activates ERK in them (7–11). Because of this delay, such feedback can generate waves of ERK activation that propagate from cell to cell across the monolayer. Such ERK waves have been implicated in collective cell migration (7–9), development (10, 12), homeostasis (10, 13, 14), and cell fate decisions (8, 13, 15, 16). However, ERK signaling is only one component of a broader mechanochemical feedback system that can also involve calcium signaling, mechanosensitive ion channels, gap junctions, integrin adhesions, and actin-based protrusions (17–22).

Mechanochemical waves have been studied mostly in flat epithelial monolayers or developing tissues, where cells form a continuous sheet and mechanical coupling occurs predominantly within the plane of the tissue. It remains unclear whether similar feedback principles exist in mesenchymal cell collectives embedded in fibrous 3D extracellular matrices. These cells face a mechanical environment that differs from epithelial monolayers: they generate strong contractile forces and remodel, stiffen, and invade the surrounding matrix (3, 5, 23–25). However, whether such 3D systems can develop coordinated mechanochemical dynamics, which signals and cytoskeletal structures coordinate these, and whether mechanical communication extends beyond individual cell collectives remain open questions.

Here, we report that fibroblasts in 3D collagen-embedded spheroids (26, 27) and engineered microtissues attached to flexible pillars (28–30) develop collective force oscillations, which propagate as calcium-dependent mechanical waves throughout spheroids and microtissues. These force oscillations are regulated by integrin ligation in actin-rich cell protrusions, ERK signaling, gap-junctional communication, and Piezo1-mediated mechanosensing. Notably, ERK signaling also controls cell invasion into the surrounding matrix, whereas high overall cellular contractility per se is not sufficient. In addition, we demonstrate that neighboring spheroids mechanically synchronize their force oscillations through the collagen matrix across millimeter distances. Together, these findings identify oscillatory mechanochemical feedback as a mechanism by which fibroblast collectives coordinate force generation and long-range mechanical communication in 3D extracellular matrices.

## Results

### Measuring Collective Cell Forces

To measure force generation in a 3D connective tissue model, we embed spheroids together with 4 µm fiducial markers in collagen type I. The spheroids are assembled from 4000 patient-derived primary human cancer-associated fibroblasts (CAFs) isolated from luminal B breast tumors. Over 24 h, we image the equatorial plane of each spheroid and measure the cell-generated matrix deformations from the displacement of the fiducial markers (Figure 1a, left). We take advantage of the spherical symmetry of this geometry and reconstruct spheroid contractile forces using collagen-specific, finite-element-based material lookup tables, as described previously (26). These tables relate local matrix deformations to contractile force based on the deformation magnitude and the distance from the spheroid center, both normalized to spheroid size (Figure 1a, right).

**Figure 1.**
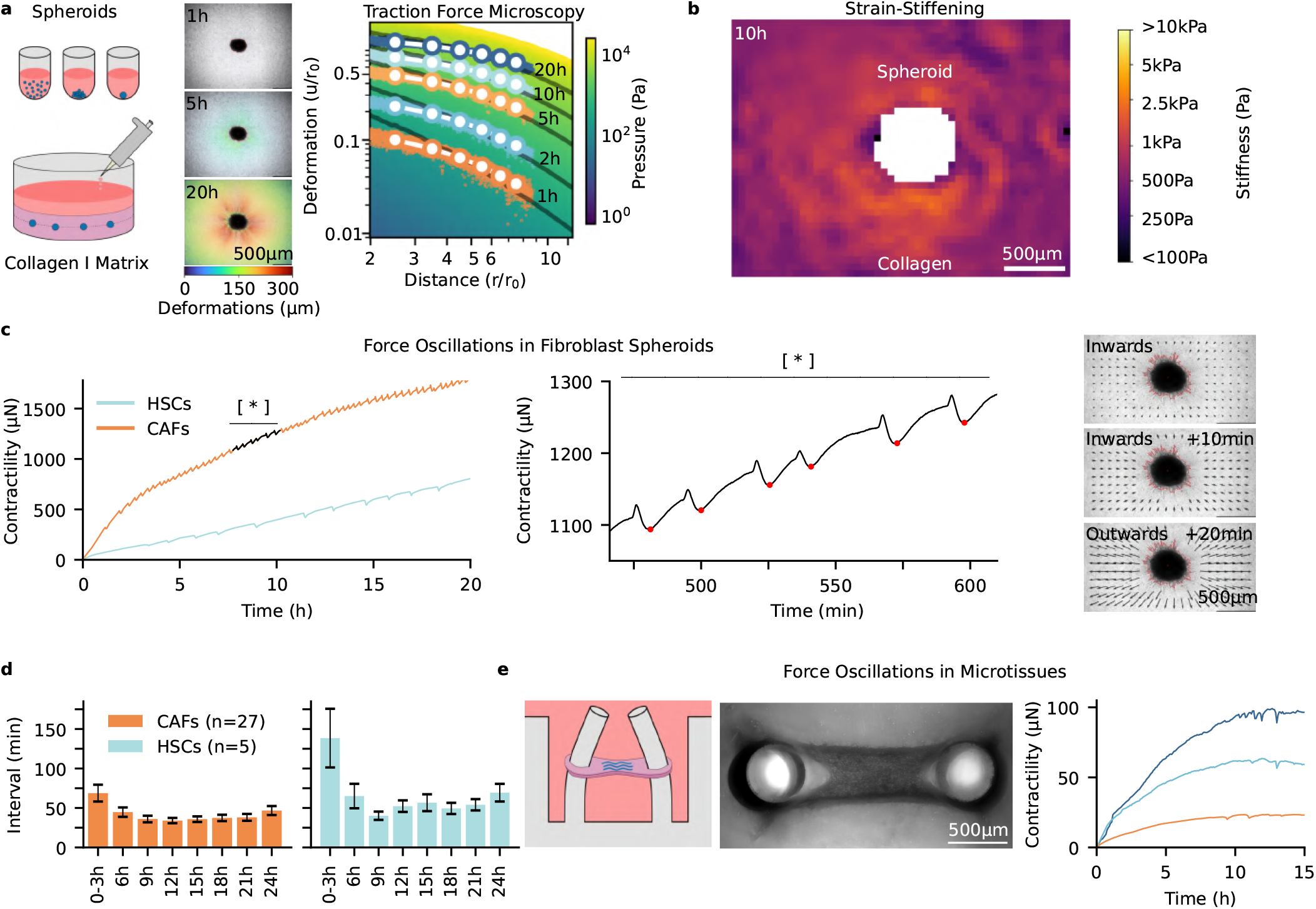
Fibroblast spheroids periodically contract 3D collagen matrices. **a**, Spheroids are generated from 4000 human CAFs cultured for 1 day in non-adhesive U-well plates (top left). The following day, the spheroids are embedded in double-layered 1.2 mg/ml collagen matrices (bottom left). Matrix deformations around the spheroids are measured dynamically from brightfield microscopy images using embedded microbeads tracked with particle image velocimetry (26, 32). Brightfield microscopy images of spheroids embedded in collagen gels are shown at different times (center). The accumulated matrix deformations are indicated by the colored arrows. Right image shows a material-specific lookup-table for reconstructing the contractile pressure (green-yellow) of a spheroid, based on the magnitude of the matrix displacements u and their distance to the spheroid center r, both normalized to the spheroid radius r0 as described in (26). Measured matrix deformations at different times are shown in different color (white dots indicate running average over distance shells). Black lines indicate the best-fit contractile pressure to the experimental data. **b**, Stiffness map of a 1.2 mg/ml collagen gel around a spheroid (white) 10 h after embedding. Stiffness values (color-coded) of the collagen gels are derived from the measured matrix deformations and the nonlinear elastic material model (26, 33, 34). **c**, Fibroblast spheroids increase contractile forces over time and generate force oscillations (left). The middle panel highlights the time interval marked by the asterisk in the left panel. Consecutive force oscillations are marked by red dots. During each oscillation, force briefly increases, then rapidly declines and slowly recovers. The right panels show the inward matrix deformation followed by outward relaxation during one oscillatory event. Similar force oscillations are observed for human breast-cancer associated (CAFs) and human liver fibroblasts (HSCs). **d**, Time interval between force oscillations of spheroids in 1.2 mg/ml collagen gels averaged over 3 hours. Bars indicate mean±se of 5 or 27 individual measurements, each containing typically 4-10 spheroids. For both CAFs and HSCs spheroids, the time interval between pulses decreases within the first 9 h and then remains relatively constant throughout the 24 h measurement period. **e**, Force oscillations also emerge in self-assembled microtissues generated from 4000 CAFs in 0.6 mg/ml collagen around two flexible pillars (28, 29). The scheme (left) and bright-field image (middle) show the experimental setup. The force time curves (right) show force oscillations in three individual microtissues (colored curves), which are also visible as oscillatory pillar bending in SI Video 6.

**Figure 2.**
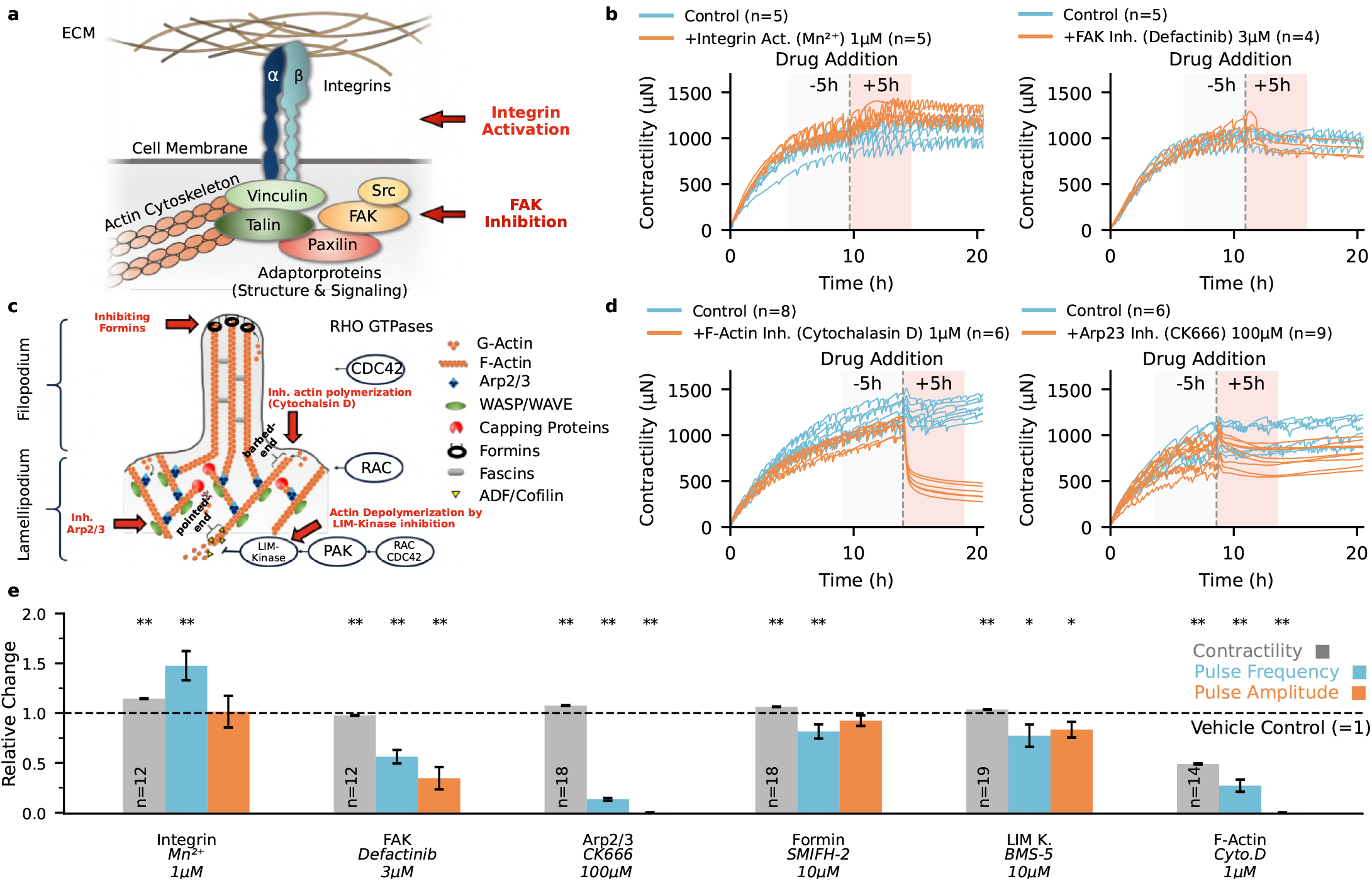
Force oscillations are mediated through integrins and actin protrusions. **a**, Schematic representation of integrin cell surface receptors (subunits *α* and *β*), which link the outer collagen matrix to the actin skeleton through various adaptor proteins. **b**, Contractile forces of spheroids are shown before and after integrin-activation using Mn^2+^ (orange, left) and focal adhesion kinase (FAK) inhibition using Defactinib (orange, right). Force curves for untreated control spheroids are shown in blue. Gray dotted line indicates the time of the drug addition, and n indicates the number of spheroids measured. **c**, Schematic representation of the actin organization and its regulation in lamellipodial and filopodial protrusions. **d**, Contractile forces of spheroids before and after cytochalasin D treatment to depolymerize F-actin (left), and before and after CK666 treatment to inhibit Arp2/3 complexes in the lamellipodium (right). **e**, Contractility (gray), pulse frequency (blue), and pulse amplitude (orange) of force oscillations in spheroids are evaluated within a 5 h window before (gray shaded area in b, d) and after drug addition (orange shaded area in b, d). Bars show the mean*±*se of the relative change after treatment, normalized to the corresponding change in control experiments (solvent addition) within the same time intervals. Target and drugs are indicated below the bars (see SI Fig. 9 for more details). Significance is determined by a paired t-test, where * indicates p<0.05 and ** indicates p<0.01. Data from individual experiments and absolute values are provided in SI Fig. 10 and 12.

### Synchronization of collective cell forces

Over time, CAF spheroids exert progressively increasing forces on the surrounding collagen, which reorganize the matrix fibers and facilitate cell invasion (Figure 1a-b, SI Fig. 1-2, SI Video 1-7, (31)). Within 2–6 h after embedding, spheroids develop collective force oscillations, indicating that force generation has become coordinated across the spheroid (Figure 1c, SI Video 1-7). The interval between oscillations is not strictly periodic but averages around 30 min, with a typical pulse amplitude of around 10 µN. Each oscillation typically starts with a brief force increase, followed by a rapid force decrease and a slower recovery (Figure 1c).

To test whether this oscillatory behavior is specific to spheroids from primary luminal B breast cancer-associated fibroblasts (CAFs) or represents a more general feature of collective force generation, we measure spheroids assembled from other primary mesenchymal cell types, including fibroblasts from healthy breast tissue (SI Video 2), cancer-associated fibroblasts from adenocarcinoma (SI Fig. 3), and hepatic stellate cells (HSCs) (Figure 1c-d, SI Video 5. We also generate microtissues from primary luminal B CAFs and cancer-associated fibroblasts from adenocarcinoma (28– 30). These microtissues self-assemble from a cell-collagen mixture between two flexible pillars and allow us to measure collective forces from pillar deflections (Figure 1e, SI Fig. 3). In all of these primary fibroblast systems, we observe similar force oscillations, suggesting a conserved mechanism for coordinated force generation in primary fibroblast collectives. This similarity extends to the timing of the oscillations: in both primary CAF and HSC spheroids, pulses typically start within the first 6 h after embedding in collagen, and their average interval (*≈*30 min) remains nearly constant over 24 h (Figure 1d). Furthermore, in both systems, the pulse amplitude and pulse frequency increase with increasing cell contractility (SI Fig. 4). We also observe weaker, less pronounced pulsing in the less contractile mouse fibroblast cell lines (SI Fig. 5), but do not detect visible pulses in a range of other cell lines, including MDA-MB-231, MCF7, HT1080, U-87, and A-172 (data partly shown in (26)). Together, these findings suggest that pronounced force oscillations are a robust feature of primary fibroblast systems but are not universally observed in established mesenchymal cell lines.

**Figure 3.**
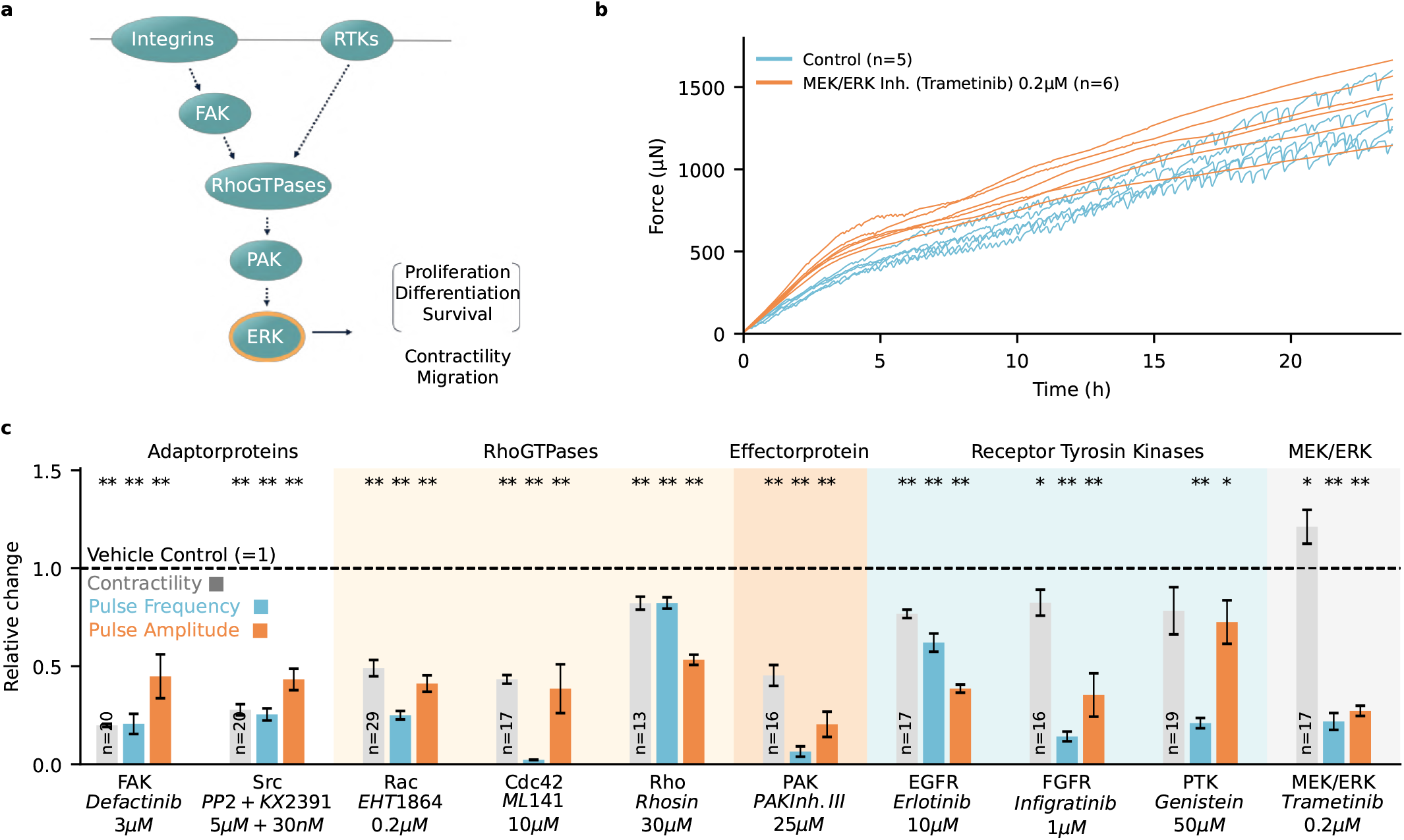
Force oscillations are mediated via the ERK signaling pathway. **a**, Schematic representation of the downstream signaling from integrins and receptor tyrosine kinases (RTKs) to extracellular signal-regulated kinase (ERK). **b**, Time development of forces in individual spheroids treated with the MEK/ERK inhibitor Trametinib (orange) and in control spheroids (blue). **c**, Small molecules are used to inhibit different signaling molecules in the ERK pathway during the full 24 h spheroid traction force measurements: focal adhesion kinase (FAK), Src kinase (Src), Rho GTPases (Rac, Cdc42 and Rho), p21-activated kinase (PAK), fibroblast growth factor receptor (FGFR), and extracellular signal-regulated kinase (ERK). Contractility (gray), pulse frequency (blue), and pulse amplitude (orange) are calculated by averaging over the 24 h measurement period. Bars represent mean*±*se of relative changes compared to sham control. Targets and drugs are indicated below the bars (see SI Fig. 9 for more details). Significance is determined by a paired t-test, where * indicates p<0.05 and ** indicates p<0.01. Raw data from individual experiments are shown in SI Fig. 11.

**Figure 4.**
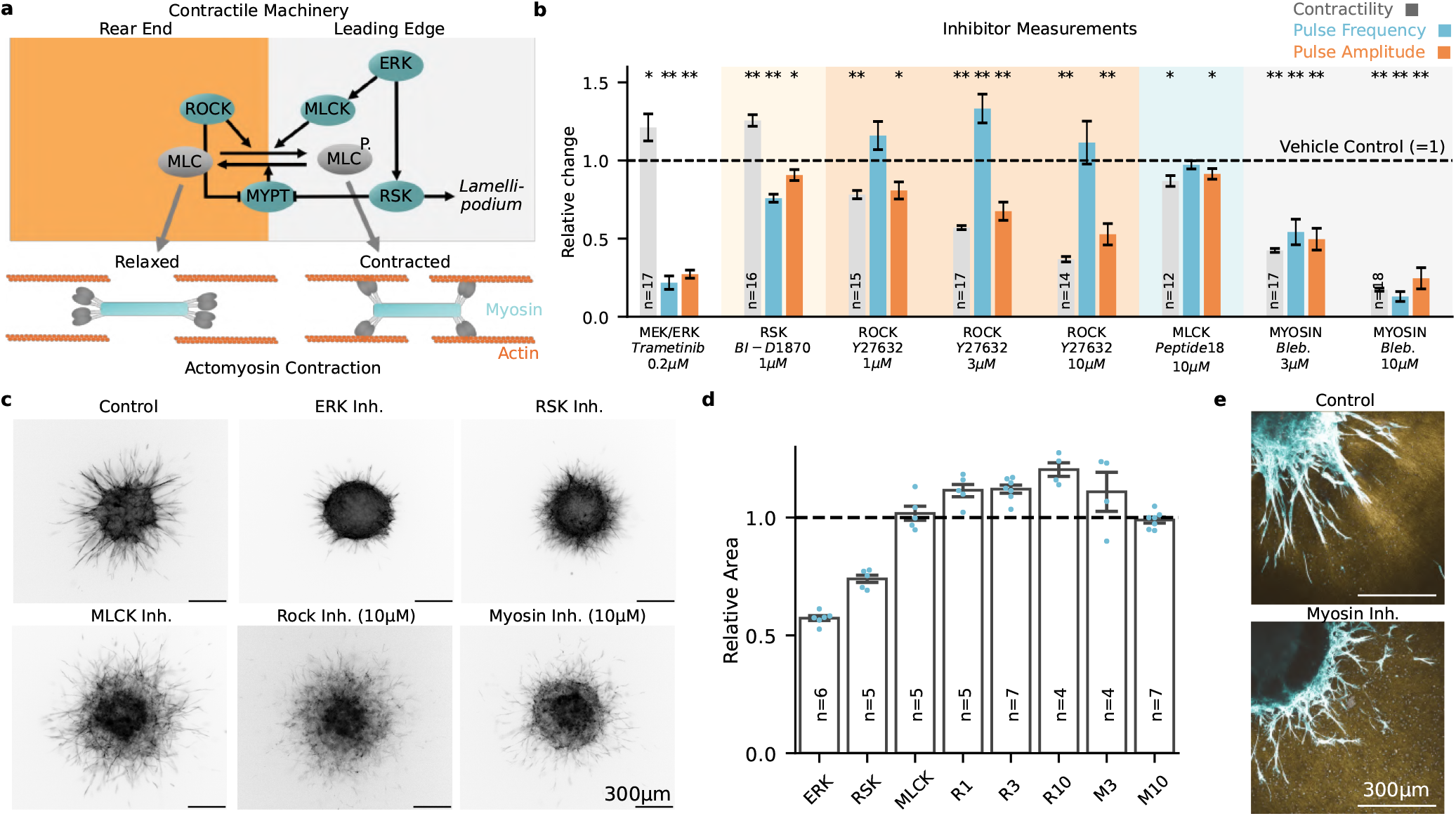
ERK-dependent force oscillations, but not overall contractility, are associated with collective cell invasion. **a**, Schematic representation of downstream signaling from ERK and rho-associated kinase (ROCK) to myosin light chain (MLC) phosphorylation regulated acto-myosin contraction adapted from (38). Signaling involves myosin light chain kinase (MLCK), myosin phosphatase (MYPT), and ribosomal s6 kinase (RSK). To investigate the downstream signaling of force oscillations, small molecules are used to inhibit different signaling molecules in this pathway during the full 24 h traction force measurements. **b**, Contractility (gray), pulse frequency (blue), and pulse amplitude (orange) are calculated by averaging drug-treated spheroids over the 24 h measurement period. Bars represent mean*±*se relative to sham control. Targets and drugs are indicated below the bars (see SI Fig. 9 for more details). Significance is determined by a paired t-test, where * indicates p<0.05 and ** indicates p<0.01. Raw data from individual spheroids are provided in SI Fig. 15. **c**, Fluorescence microscopy images showing the actin-stained cells at the spheroid equatorial plane after 24 h in 1.2 mg/ml collagen gels for drug-treated and control spheroids. **d**, As a measure of cell invasion into the collagen matrix, the relative changes in the actin-stained cell area between drug-treated and control spheroids after 24 h are calculated for ERK-, RSK-, and MLCK inhibition as well as for Myosin (M)- and ROCK (R) inhibition at different doses. **e**, Overlay of confocal microscopy images of the actin-stained cell area (blue) and confocal reflectance microscopy images of the collagen fiber structure (brown) after 24 h, showing increased alignment of cells and collagen fibers around control spheroids compared to myosin-inhibited spheroids, consistent with previously observed correlation between cell contractility and matrix fiber alignment (31).

**Figure 5.**
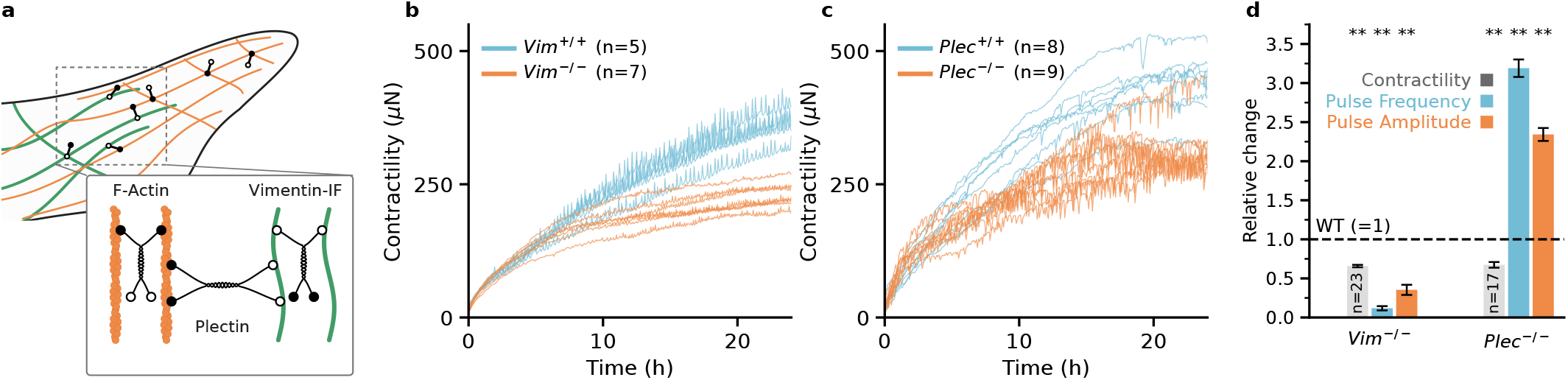
The intermediate filament system modulates force oscillations. **a**, Schematic representation of plectin-mediated crosslinking of F-actin (orange) and vimentin intermediate filaments (green). The upper schematic shows plectin linkers (black) as polar complexes with N-terminal actin-binding (filled circle) and C-terminal intermediate-filament-binding (open circle) regions connected by a rod-like spacer. The inset shows the proposed structural organisation of a parallel plectin dimer (39). In this model, each circle in the upper schematic represents a pair of terminal regions that may bind two nearby filaments of the same type. **b**, Time development of forces over 24 h in individual spheroids formed from wild-type (blue) or vimentin-deficient (*Vim*^*−/−*^; left) or plectin-deficient (*Plec*^*−/−*^; right) mouse fibroblasts (orange). Each curve represents one spheroid. **c**, Relative contractility at 24 h (gray), pulse frequency (blue), and pulse amplitude (orange) in vimentin- and plectin-deficient spheroids. Values are normalized to the respective wild-type mean within each experiment. Bars represent mean *±* se across spheroids from three independent experiments for vimentin and two for plectin. Numbers inside the bars indicate the number of knockout spheroids per group. Significance is determined by an student t-test, where *∗*indicates *p <* 0.05 and *∗∗* indicates *p <* 0.01.

### Force oscillations depend on the mechanical environment

Given the robustness of this behavior across different fibroblast subtypes, we investigate whether force oscillations are driven by mechanical coupling between cells through their environment. In the experiments reported above, we have embedded the cells in 1.2 mg/ml collagen. Although the collagen matrix at this concentration is soft (Young’s modulus *≈*200 Pa), it exhibits non-linear strain-stiffening at higher strains above *≈*5 *−*10% (SI Fig. 6 (35, 36)), leading to a strong increase in matrix stiffness over time as the spheroids contract (Figure 1b, SI Video 8).

**Figure 6.**
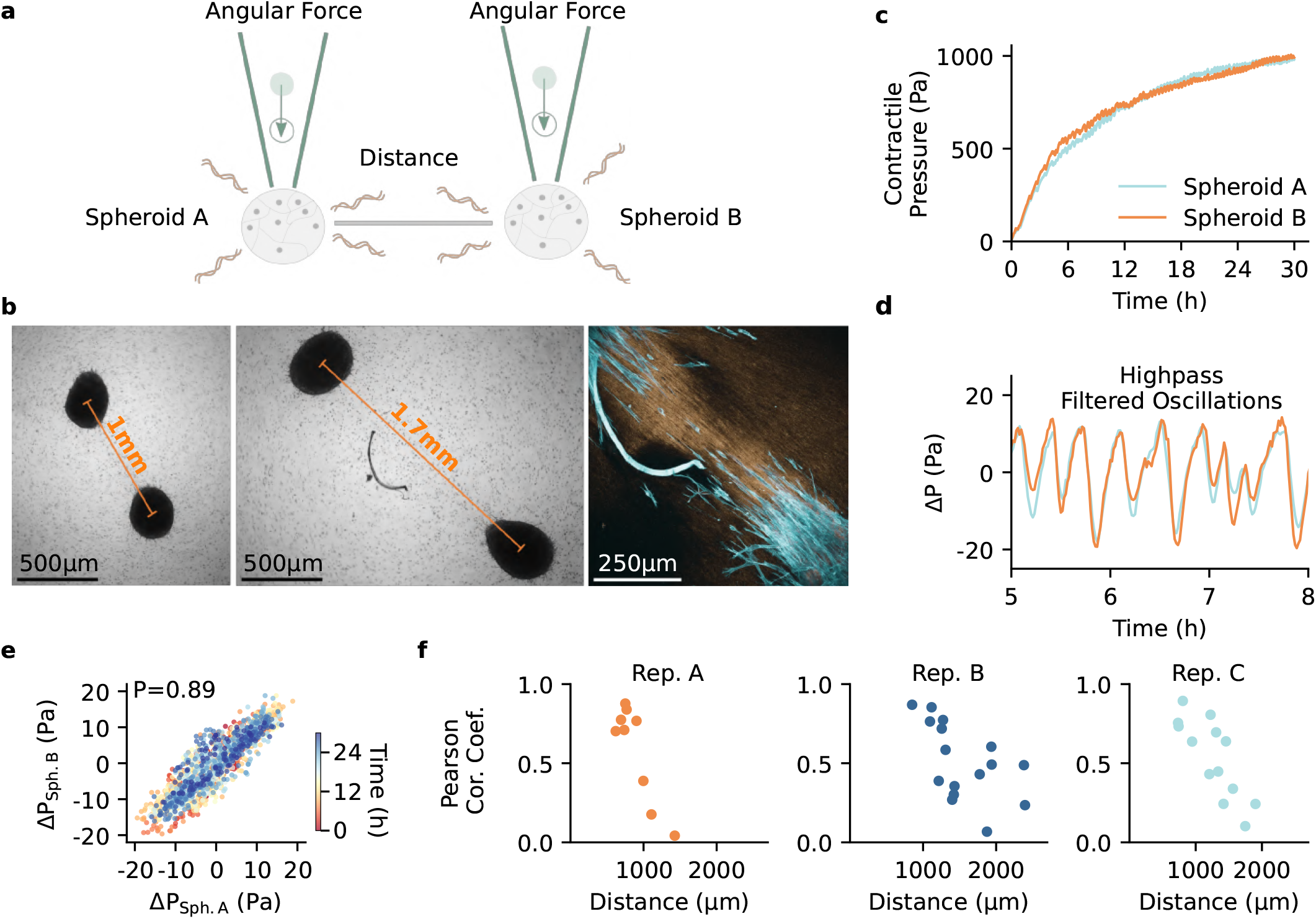
Mechanical synchronization of collective forces in 3D. **a**, Schematic of the experimental setup: CAF-spheroids are embedded close to each other in 3D 1.2 mg/ml collagen gels. **b**, Spheroid pairs at varying distances within collagen gels containing 4 µm-sized microparticles as fiducial markers. Left and middle: Brightfield images after 3 h. Right: Confocal image after 24 h showing collagen (brown) and actin-stained cells (blue). Increased fiber alignment between neighboring spheroids was previously quantified in SI Fig. 12 of (31) (see SI Fig. 2). **c**, The contractile pressure of individual spheroids is measured over 1 day based on the matrix deformation within a 5°angular cone that is directed perpendicular to a line connecting adjacent spheroids. **d**, The pressure curves of each spheroid are detrended using a Butterworth high-pass filter (48) with a cutoff frequency of 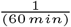 (right). **e**, The high-pass filtered pressure curves of adjacent spheroids are cross-correlated, and the Pearson’s correlation coefficient (P) is calculated as a measure of the synchrony between the curves. To visualize the correlation, the pressure curves of two adjacent spheroids (SI Video 13) are plotted against each other, where the color (red-blue) encodes the measurement time. **f**, Pearson’s correlation coefficient as a function of the distance between adjacent spheroids. Results from three independent experiments are shown, each showing a positive correlation and hence force synchronization between spheroids for distances of up to 2 mm.

To study whether feedback from the non-linear strain-stiffening of collagen promotes force oscillations, we embed spheroids in matrices with varying degrees of strain-stiffening. First, we test Matrigel, a basal membrane extract, which exhibits comparable stiffness to collagen but a much less pronounced strain-stiffening, starting at 40-50% strain (SI Fig. 6). In Matrigel, we observe reduced force oscillations (SI Video 9). Similarly, in purely linear elastic materials such as Gelatin Methacryloyl (GelMA) or alginate dialdehyde (ADA) gels, which lack strain-stiffening (SI Fig. 6), the force oscillations are strongly reduced (SI Video 9, 10).

To investigate whether strain-stiffening, rather than the baseline stiffness, drives this behavior, we compare differently concentrated collagen gels. Despite their different baseline stiffness, these gels all share similar degrees of strain-stiffening ((35), SI Fig. 6) and plasticity under repetitive deformation (SI Fig. 7). We observe a similar pulse frequency across all concentrations with slightly increased frequencies in the lower-concentrated gels (SI Fig. 8).

**Figure 7.**
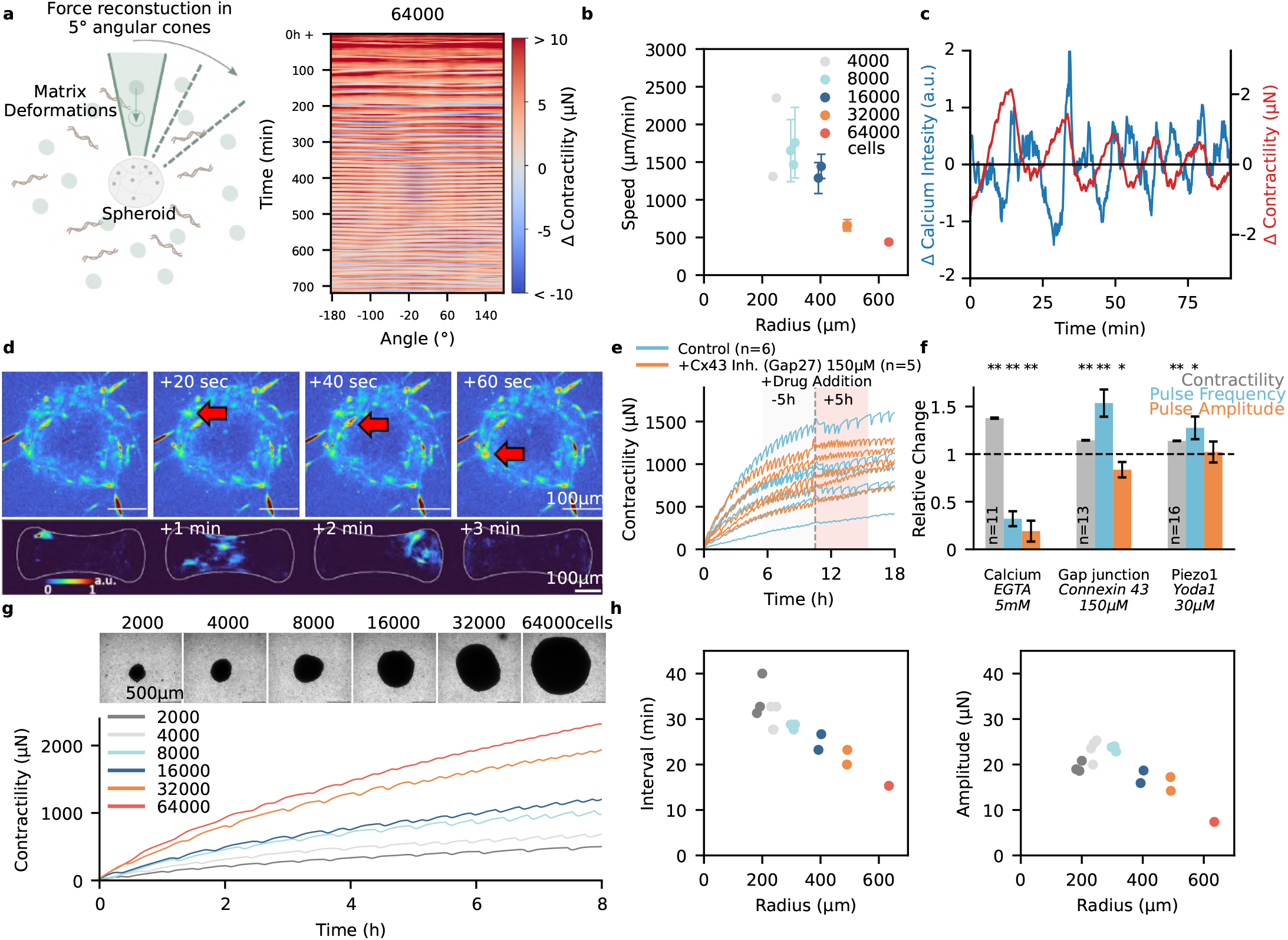
Cell-cell communication is mediated by mechanical and calcium waves. **a**, Angular spheroid forces are reconstructed from matrix deformations within 5° angular sectors around the spheroid over time (left). The kymograph shows changes in force between consecutive 1-min time points for the different angular sectors of a spheroid generated from 64000 cells (right). Kymographs for spheroids of different sizes are shown in SI Fig. 18. **b**, Estimated propagation speeds of mechanical waves in spheroids of different sizes. Points show the mean±se wave speed across pulses for each spheroid (see SI Fig. 22 for more details). Colors indicate the initial cell number. **c**, Spheroid contractility (red) and intracellular calcium intensity (blue) during force oscillations. Contractility and calcium signals are measured in the spheroid equatorial plane by confocal reflection and confocal fluorescence imaging, respectively. Cells are loaded with 4 µM Fluo-8 AM. **d**, Pseudocolored images of intracellular calcium in a human breast cancer-associated fibroblast spheroid embedded in a 1.2 mg/ml collagen gel (top; 4 µM Fluo-8 AM) and in a microtissue generated from adenocarcinoma-associated fibroblasts (bottom; 2 µM Fluo-4). The images show calcium waves propagating around the spheroid and across the microtissue over time (SI Video 19 and 20). **e**, Contractility of spheroids treated with the connexin-43 mimetic peptide Gap27 (150 µM; orange) and corresponding control spheroids (blue). The vertical gray dotted line marks drug addition, and n indicates the number of spheroids. **f**, Relative changes in contractility (gray), pulse frequency (blue), and pulse amplitude (orange) after extracellular calcium chelation with EGTA (5 mM), gap junction inhibition with Gap27 (150 µM), or Piezo1 activation with Yoda1 (30 µM). The parameters are evaluated within 5 h windows before and after EGTA or Gap27 addition and within 3 h windows before and after Yoda1 addition. Bars show the mean±se of the relative change after treatment, normalized to the corresponding change in control experiments over the same time intervals. Targets and compounds are indicated below the bars (see SI Table 9). Significance is determined using a paired t-test, where * indicates p < 0.05 and ** indicates p < 0.01. Results from individual experiments and absolute values are shown in SI Fig. 21. **g**, Representative brightfield images and contractility traces of spheroids generated from 2000, 4000, 8000, 16000, 32000, or 64000 cells. **h**, Overall contractility, pulse amplitude, and pulse interval as functions of spheroid radius, evaluated during the first 12 h after embedding.

### Integrin-mediated mechanotransduction in actin protrusions

Given our finding that matrix mechanical properties modulate force oscillations, we next test whether focal adhesions mediate this mechanical sensitivity (Figure 2a). For this, we target different signaling and structural molecules by adding activating or inhibiting drugs to the culture medium and compare the total static force, pulse frequency, and pulse amplitude in a 5 h window before and after drug addition (SI Fig. 9).

First, we focus on integrins, which directly connect the cells to the surrounding matrix (Figure 2a). Different doses of the integrin-activating ion manganese show that at high levels of activation, the spheroids respond with an increase in static force (SI Fig. 10), whereas at low levels of activation, they respond with an increased frequency of force oscillations (Figure 2b,e, SI Fig. 10). Integrin engagement activates intracellular signaling through focal-adhesion-associated proteins, including FAK and Src. Consistent with this, inhibition of either kinase reduces both the frequency and amplitude of the force oscillations (Figure 2b, e, SI Fig. 11). As integrins are mechanically connected to the cytoskeleton, we next perturb the actin network (Figure 2c). F-actin depolymerization strongly reduces both static force and pulse frequency (Figure 2d-e, SI Fig. 12). The reduction in force is expected because actomyosin contractility is required for force generation, whereas the lower pulse frequency indicates that an intact actin network is also required for the oscillatory dynamics.

Since F-actin-based cell protrusions are critical for mechanosensing processes in individual cells (37), we next examine the actin-regulatory pathways associated with lamellipodia and filopodia. Inhibition of Arp2/3-mediated branched actin nucleation in lamellipodia using CK666 strongly reduces pulse frequency while leaving static force largely unchanged (Figure 2d-e, SI Fig. 12). The addition of SMIFH-2 to inhibit formin-mediated actin-nucleation in filopodia produces a similar but less pronounced effect (Figure 2e, SI Fig. 12). Consistent with a role for actin-based cell protrusion in mediating mechano-responsive force oscillations, the inhibition of the ADF/cofilin-regulator LIM kinase with BMS-5 also reduces pulse frequency and pulse amplitude (Figure 2e, SI Fig. 12).

Together, these experiments identify focal-adhesion signaling and actin-rich protrusions as key regulators of collective force oscillations.

### ERK-mediated mechanotransduction

To investigate how the signals initiated at the cell–matrix interface propagate further through intracellular pathways over longer timescales, we extend our drug screening to a 24 h observation period. During this period, we quantify the overall force, pulse frequency, pulse amplitude, and cell invasion into the surrounding matrix.

Since integrins, jointly with receptor tyrosine kinases (RTKs), regulate actin cytoskeleton reorganization through common downstream signaling pathways including Rho-family small GTPases (Figure 3a), we first focus on RTKs.

We inhibit various RTKs, including EGFR, FGFR, and VEGFR, using the specific inhibitors erlotinib, infigratinib, and axitinib, respectively, as well as the broad protein tyrosine kinase (PTK) inhibitor genistein. All treatments reduce pulse frequency, while the overall force is only slightly reduced (Figure 3c, SI Fig. 13, 14). This observation aligns with the established crosstalk between integrins and RTKs (19, 20).

Downstream of integrins and RTKs, we examine the small GTPases Rac, Cdc42, and Rho (Figure 3c), which regulate lamellipodia, filopodia, and stress fiber dynamics, respectively (21, 22). Consistent with the role of actin-rich protrusions in force oscillations, inhibition of Rac or Cdc42, as well as of their downstream effector p21-activated kinase (PAK), strongly decreases pulse frequency (Figure 3c, SI Fig. 11). Notably, inhibition of Rho has only a minor effect on pulse frequency (Figure 3c, SI Fig. 11), even though it reduces pulse amplitude and overall forces. These findings further highlight the role of actin-rich protrusions in regulating force oscillations.

Integrin and RTK signaling converge on the extracellular signal-regulated kinase (ERK) pathway (Figure 3a-b). Previous studies have shown that ERK-mediated mechanochemical feedback regulates cell contractility and migration (7– 9). Consistent with these findings, inhibition of mitogen-activated protein kinase 1 and 2 (MEK1/2), the kinases that activate ERK1/2, with trametinib strongly reduces the frequency and the amplitude of force oscillations in our 3D fibroblast spheroids, while overall spheroid contractility remains high (Figure 3b-c, SI Fig. 11). These findings identify MEK–ERK signaling as a key regulator of force oscillations in 3D fibroblast spheroids.

### Contractile machinery and matrix invasion

We further examine the contribution of the actomyosin contractile machinery (Figure 4a). Across spheroids, higher overall contractility is associated with higher pulse frequency (Figure 1c, SI Fig. 4). In line with this, inhibition of non-muscle myosin II with blebbistatin reduces both overall contractility and pulse frequency (Figure 4b).

Actomyosin contractility is regulated by several signaling pathways involving Rho-associated kinase (ROCK) and myosin light-chain kinase (MLCK) (Figure 4a). Consistent with our observation that Rho-inhibition does not strongly affect pulse frequency, inhibition of ROCK also reduces overall contractility but does not reduce pulse frequency (Figure 4b, SI Fig. 15-16). Similarly, the inhibition of MLCK does not strongly affect pulse frequency (Figure 4b, SI Fig. 15-16).

By contrast, inhibition of p90 ribosomal S6 kinase (RSK), which acts downstream of ERK on protrusive rather than cortical actomyosin (38), alters the pulse frequency (Figure 4b, SI Fig. 15). The direction of change, however, depends on the time point of drug addition after embedding the spheroid. When the inhibitor is added at the beginning, as in the experiments described above, the frequency of force oscillations is reduced (Figure 4b, SI Fig. 15). When the inhibitor is added after 10 h, by which time actin-rich protrusions are fully developed, the frequency of force oscillations is increased. In both cases, RSK inhibition increases static forces (SI Fig. 16). These findings highlight that force oscillations are not simply determined by the magnitude of contractility but rely on additional signaling in actin-rich protrusions.

We then investigate how inhibition of the contractile machinery affects cell invasion into the surrounding collagen matrix after 24 h. Despite strongly reduced contractility, inhibition of ROCK or non-muscle myosin II does not prevent invasion into 1.2 mg/ml collagen gels (Figure 4c-e, SI Video 11). Instead, these perturbations produce a less compact, more dispersed invasion pattern and reduce collagen-fiber alignment. By contrast, inhibition of ERK or RSK significantly reduces cell invasion despite sustained high contractility (Figure 4c-d, SI Fig. 17, SI Video 12). These findings indicate that ERK–RSK signaling and the resulting contractile oscillations, rather than static contractility alone, are required for efficient invasion in our 3D system.

### Intermediate filaments modulate force oscillations

After identifying a role for integrin–actin–ERK signaling in regulating force oscillations, we next turn to vimentin, the major intermediate filament protein in fibroblasts (Figure 5a). Vimentin binds ERK directly and stabilizes its activation (40). Like actin, it also forms a mechanically supportive network that contributes to focal adhesion organization, force transmission, and dissipation (41).

We find that vimentin-deficient spheroids show reduced contractility, pulse frequency, and pulse amplitude (Figure 5b,d). The reduced ERK phosphorylation reported in vimentin-deficient cells may contribute to the lower pulse frequency and amplitude observed here (40). In addition, vimentin loss has been reported to cause RhoA/ROCK hyperactivation and increased stress-fiber assembly (42), consistent with our finding that Rho or ROCK inhibition had only minor effects on pulse frequency.

The vimentin and actin networks are interconnected by cytolinkers such as plectin (Figure 5a). Plectin crosslinks vimentin filaments (43), connects the vimentin network to actin and integrin-based adhesions (44, 45), and directly crosslinks actin filaments (46).

We find that plectin-deficient spheroids also show reduced overall contractility, but their pulse frequency and amplitude are increased (Figure 5c,d). These cells are known to exhibit faster actin turnover (47), which may contribute to the enhanced oscillations, given our finding that actin-rich protrusions rather than Rho/ROCK signaling regulate pulse frequency.

Together, these findings suggest that vimentin supports oscillatory force transmission, whereas plectin-mediated cytoskeletal coupling limits force oscillations, potentially by constraining actin dynamics.

### Mechanical communication through 3D collagen

To test whether force oscillations can synchronize between spatially separated spheroids through the collagen matrix, we embed spheroids at varying distances from one another. For each spheroid, we reconstruct the contractile pressure from matrix deformations within angular sectors oriented perpendicular to the line connecting the spheroid pair (Figure 6a-c). We then high-pass filter the resulting pressure curves to isolate the oscillatory component (Figure 6d) and quantify synchrony between neighboring spheroids using the Pearson correlation coefficient (Figure 6e).

Our data show that force oscillations of neighboring spheroids become synchronized, with positive correlations observed over distances of up to 2 mm (Figure 6f, SI Video 13). We also observe strong alignment of collagen fibers between neighboring spheroids, consistent with the formation of mechanically coupled matrix regions, as previously characterized (SI Fig. 2, (31)). Together, these findings indicate long-range mechanical coupling between spheroids through the 3D collagen matrix.

### Calcium signaling and cell-cell communication

We next examine how neighboring cells within a spheroid communicate. Previous studies have shown that coupled mechanical and biochemical activation waves propagate between neighboring cells in 2D epithelial monolayers (7, 8). In our 3D system, strong light scattering and limited spatial resolution prevent us from resolving individual cells within the spheroid. However, by reconstructing contractile pressure in angular sectors around the spheroid, we detect mechanical waves that propagate around the spheroid during individual force pulses at speeds of several hundred micrometers per minute, highlighting a mechanical signal transmission between neighboring cells (Figure 7a-b, SI Fig. 18, SI Video 14–17).

We further find that larger spheroids generate greater contractile forces and exhibit higher pulse frequencies during the first 12 h (Figure 7g,h). At these early time points, the pulse amplitude initially increases with spheroid size but decreases in larger spheroids after 12 h (SI Fig. 19). This reduction may reflect insufficient mechanical recovery between the more frequent pulses, or the initiation of multiple mechanical waves at different sites, rather than a single wave propagating around the spheroid. Such waves may partially overlap and produce irregular, spatially heterogeneous contractions, visible as wobbling motions in the timelapse recordings and as overlapping patterns in the kymographs (SI Video 14–17, SI Fig. 18). Force oscillations can also pause for several hours before resuming. During these intervals, the kymographs show traveling-wave activity rather than coherent spheroid-wide pulses (SI Fig. 19, SI Fig. 18, SI Video 18).

Since mechanical force generation is closely linked to calcium signaling, we simultaneously monitor intracellular calcium and force oscillations. Confocal fluorescence imaging shows calcium waves that propagate at several hundred µm/min around the spheroid or across a microtissue at the onset of a force pulse (Figure 7c-d, SI Fig. 20, SI Video 19-20).

Chelation of extracellular calcium with EGTA reduces both pulse frequency and amplitude, consistent with a role for calcium signaling in force oscillations (Figure 7f, SI Fig. 21). By contrast, inhibition of gap junction communication with Gap27 increases pulse frequency while reducing pulse amplitude (Figure 7f, SI Fig. 21). This response is consistent with reduced intercellular synchronization. Accordingly, local contractile events may occur more independently and therefore more frequently, but combine less coherently into large spheroid-scale force pulses.

We then examine the contribution of mechanosensitive calcium channels. Activation of Piezo1 with Yoda1 increases pulse frequency (Figure 7f, SI Fig. 21). Likewise, inhibiting mechanosensitive channels with GsMTx-4 reduces pulse frequency and pulse amplitude (SI Fig. 14). Together, these findings indicate that calcium signaling and mechanosensitive channel activity regulate collective force dynamics.

## Discussion

Here, we show that fibroblast collectives coordinate force generation through oscillatory mechanochemical feedback involving cell-matrix coupling, intracellular signaling, and intercellular communication. Such coordination is relevant to multicellular processes such as tissue remodeling, wound healing, and cancer invasion.

Using multicellular 3D spheroids and engineered microtissues, we find that collective force generation occurs in oscillations with a characteristic interval of approximately 30 min. These oscillations are regulated by integrin and receptor tyrosine kinase signaling through Rho-family GTPases, PAK, and the MEK-ERK pathway. Within spheroids, mechanical and calcium waves propagate between neighboring cells, whereas oscillations of separate spheroids synchronize through the collagen matrix over distances of up to 2 mm.

Our findings in 3D fibroblast systems extend previous observations of ERK-mediated mechanochemical waves in 2D epithelial monolayers (7, 8, 13) and developing tissues (10, 11). Such waves have been implicated in collective cell migration (7–9), morphogenesis and development (10–12), and tissue homeostasis (13, 14).

In this work, we observe ERK-dependent force oscillations in fibroblasts from different tissue origins, both within and between 3D cell collectives. In fibroblasts, ERK signaling and force oscillations may be of particular interest, since both are known to play an important role in the context of fibrosis and are explored as a therapeutic target (49–51). In 2D epithelial cell sheets, the mechanism that connects mechanical stretch and ERK activation has been identified (7, 8, 52). Since we also observe mechanical waves propagating between neighboring fibroblasts, a similar mechanism likely underlies the force oscillations in our 3D system. However, we have not demonstrated this in 3D, which would require sophisticated live imaging of ERK activity, for example by using biosensors and FRET imaging.

In our system, inhibition of ERK signaling reduces force oscillations and strongly impairs invasion into the surrounding collagen matrix. By contrast, reducing contractility through inhibition of ROCK or non-muscle myosin II does not prevent invasion, but changes it from strand-like collective migration along aligned collagen fibers to a less compact, more dispersed pattern (SI Fig. 17, SI Video 11). ERK inhibition, however, strongly suppresses invasion despite sustained spheroid contractility and collagen-fiber alignment (31) (SI Fig. 17, SI Video 12). These findings indicate that force oscillations and invasion share a dependence on ERK signaling. ERK signaling, rather than high overall contractility, is required for efficient collective invasion in our 3D system.

For spheroids of different sizes, we find that the estimated mechanical wave speed decreases with increasing spheroid radius (Figure 7). This size dependence coincides with a slight decrease in cell packing density for larger spheroid (SI Fig. 22), consistent with previous observations (53). Reduced packing density may weaken the mechanical coupling between neighboring cells and could possibly contribute to the slower wave propagation in larger spheroids.

Our findings identify calcium signaling as a potential link between mechanosensation, ERK activity, and the coordination of collective force generation. In our experiments, calcium waves accompany the onset of force pulses in both spheroids and microtissues (Figure 7c,d), and perturbation of gap junction communication changes pulse frequency and amplitude (Figure 7f). These observations are consistent with altered coordination of local contractile events across the fibroblast collective. Additionally, calcium-dependent N-cadherin adhesion may contribute to mechanochemical coupling between fibroblasts. The EGTA-induced increase in contractility and decrease in pulse frequency and amplitude (Figure 7f) could reflect impaired cell-cell adhesion in addition to altered calcium signaling (54). We further find that Piezo1 activation increases pulse frequency, while inhibition of mechanosensitive channels perturbs the oscillatory dynamics (Figure 7f, SI Fig. 14). Together with previous reports linking Piezo1 activity to EGFR and ERK signaling (17, 18), our results support a model in which calcium-dependent intercellular communication and mechanosensitive channel activity interact with ERK signaling to regulate collective force oscillations.

Several additional observations support a partial dissociation between overall contractility and oscillatory dynamics. Treatment with the broad-spectrum metalloprotease inhibitor GM6001 or the ADAM17/TACE inhibitor TAPI-I reduces overall contractility by *≈*30% and *≈*45%, respectively. However, GM6001 increases pulse frequency, whereas TAPI-I does not (SI Fig. 13-14). ROCK inhibition and nutrient starvation reduce contractility without suppressing pulse frequency, providing further evidence that oscillatory dynamics are not determined solely by the magnitude of force generation (Figure 4b, SI Fig. 14). Pulse frequency also increases with increasing cell passage number and prolonged time in culture, whereas overall contractility shows no corresponding increase (SI Fig. 4, 23, 24). This culture dependence suggests that prolonged culture selectively enhances the signaling pathways controlling pulse generation, potentially through increased ERK pathway activity.

Apoptosis-triggered ERK/Akt waves have been linked to epithelial survival signaling (13). In our system, force oscillations are strongly reduced by MEK/ERK inhibition, whereas AKT inhibition has little effect (SI Fig. 13-14) and apoptosis induction rather suppresses force oscillations (SI Fig. 25). This suggests that force oscillations are primarily associated with ERK signaling rather than with AKT-mediated survival signaling.

Although our data identify ERK and calcium signaling as important regulators of force oscillations, our perturbation experiments suggest that these pathways act within a broad network of signaling molecules and cytoskeletal components. Interactions within this broad network can therefore affect force oscillations indirectly. For example, hypotonic shock increases pulse frequency, whereas ethanol decreases it in our experiments (SI Fig. 13 and 14). Because hypotonicity and ethanol have previously been reported to activate and suppress ERK signaling, respectively (55, 56), these responses are consistent with ERK-dependent regulation, although ERK activity was not measured directly under these conditions.

The mechanical environment provides an additional component of this feedback. Force oscillations are pronounced in strain-stiffening collagen but are strongly reduced in Matrigel and in the linear elastic GelMA and ADA-GEL matrices (SI Video 9-10). As spheroids contract the collagen network, the matrix stiffens locally over time (Figure 1b, SI Video 8), which may progressively strengthen force transmission and integrin-dependent mechanochemical feedback. Molecular-clutch models provide a possible framework for this interpretation, as nonlinear force transmission through integrin adhesions can couple changes in matrix mechanics to intracellular mechanotransduction (57). Our experiments do not identify a single optimal matrix stiffness, but instead suggest that progressive local strain stiffening modulates this force-transmission feedback. YAP signaling represents one potential downstream component of this coupling, and its perturbation also alters pulse frequency (SI Fig. 13-14). Together, our observations support a model in which matrix stiffening alters integrin-mediated force transmission and downstream signaling, thereby enabling fibroblast collectives to remodel and dynamically sense their mechanical environment through force oscillations.

In summary, fibroblast spheroids and microtissues generate collective force oscillations through mechanochemical feedback involving integrin-dependent cell-matrix coupling, actin-rich protrusions, Rho-family GTPases, MEK-ERK signaling, calcium signaling, and mechanosensitive channels. Mechanical and calcium waves propagate within the cell collectives, while force oscillations synchronize between separate spheroids through the 3D collagen matrix over distances of up to 2 mm. ERK signaling is required not only for the oscillations but also for efficient collective invasion. Together, these findings show that mechanochemical feedback coordinates force generation, intercellular communication, matrix stiffness sensing, and invasion in 3D fibroblast collectives. Such coordination may contribute to tissue remodeling, wound healing, and cancer invasion.

## Material and Methods

### Cell culture

#### Breast cancer-associated fibroblasts (CAFs)

Primary cultures of human breast cancer-associated fibroblasts (luminal b) are isolated as previously described in (26). Cells are cultured in Epicult basal media supplemented with Supplement C (Epicult-C human media kit, Stem Cell Technologies, Vancouver), 5% FCS, 1% penicillin, and 1% streptomycin. Cell culture flasks are coated with a collagen solution (0.1 mg/ml collagen R (Matrix Bioscience, Mörlenbach) in PBS).

#### Hepatic stellate cells (HSCs)

Human primary hepatic stellate cells (HSC0141, purchased from Lonza, Basel) are cultured in Hepatic stellate growth medium (MCST250, Lonza, Basel).

#### NIH3T3 cell line

NIH3T3 mouse fibroblasts are cultured in Dulbecco’s Modified Eagle Medium (DMEM, 4.5 g/l Glucose, Thermo Fisher, Waltham) with 10% FCS, 1% penicillin, and 1% streptomycin.

#### Mouse fibroblasts

Immortalized mouse skin fibroblasts are derived from wild-type (*Plec*^+*/*+^) and plectin-deficient (*Plec*^*−/−*^) p53-deficient (*p53* ^*−/−*^) mice following protocols described previously (44, 58–60). Immortalized mouse embryonic fibroblasts (MEFs) are derived from vimentin wild-type (*Vim*^+*/*+^) and vimentin-deficient (*Vim*^*−/−*^) mice, as described previously (40), and are kindly provided by J. Ivaska (University of Turku, Finland) and J. Eriksson (Åbo Akademi University, Turku, Finland). All four cell lines are cultured in Dulbecco’s Modified Eagle Medium (DMEM, 4.5 g/l Glucose, Thermo Fisher, Waltham) supplemented with 10 % FCS, 1 % penicillin-streptomycin, and 1 % L-glutamine.

#### Cancer-associated fibroblasts from adenocarcinoma

Human primary cancer-associated fibroblasts (CAF) are kindly provided by D. Matic Vignjevic (I. Curie, Paris). These cells are isolated from a Lieberkühnian adenocarcinoma of the upper rectum and immortalized as described earlier (61) at Institut Curie Hospital, Paris, with the patient’s written consent and the approval of the local ethics committee. CAFs are used at fewer than 15 passages in culture medium composed of Dulbecco’s modified Eagle’s medium (DMEM, Thermo Fisher, Waltham) supplemented with 10% fetal bovine serum (FBS, Thermo Fisher, Waltham), 1% Insulin-Transferrin-Selenium (ITS, Sigma-Aldrich, St. Louis), 100 U/ml of penicillin and 100 µg/ml of streptomycin (Thermo Fisher, Waltham).

All cell types are kept at 37°C in an atmosphere saturated in humidity and containing 5% CO_2_.

### Hydrogel preparation

#### Collagen hydrogels

Collagen type I hydrogels are prepared from acid-dissolved rat tail (R) and bovine skin (G1) collagen (Matrix Bioscience, Germany). Both collagen types are mixed on ice at an R:G1 mass ratio of 1:2 and diluted in a solution consisting of one volume of NaHCO_3_, one volume of 10 × DMEM, and eight volumes of H_2_O. The final collagen concentrations are adjusted to 0.6 mg/ml, 1.2 mg/ml, or 2.4 mg/ml. The pH is adjusted to 9 using NaOH. Collagen gels are polymerized at 37°C, 95% humidity, and 5% CO_2_.

#### Matrigel hydrogels

Matrigel hydrogels (Corning, New York) are prepared at a final protein concentration of 10 mg/ml and polymerized at 37°C, 95% humidity, and 5% CO_2_.

#### GelMA hydrogels

Gelatin methacryloyl hydrogels (GelMA) are prepared by dissolving lyophilized GelMA at 10% and LAP photoinitiator at 0.25% in PBS (Cellink, Gothenburg). The gels are photocrosslinked using a UV lamp positioned at a distance of 15 cm for 60 seconds. This formulation is referred to as GelMA (High). A second, softer formulation, GelMA (Low), is prepared by diluting the GelMA/LAP precursor solution with PBS at a 1:2 ratio. These gels are photocrosslinked using the same UV lamp at a distance of 15 cm for 120 seconds.

#### ADA-Gel hydrogels

Alginate dialdehyde (ADA)-gelatin (ADA-GEL) hydrogels are prepared as described previously (62, 63). ADA and gelatin are separately dissolved at 5% (w/v) in PBS, sterile-filtered, and then combined to form ADA-GEL at a final polymer concentration of 2.5% (w/v). The ADA-GEL is stirred for 10 min at 37°C to obtain a homogeneous solution. After casting, the solution is placed at 4°C for 10 min and subsequently overlaid with a 0.1 M CaCl_2_ solution for 15 minutes to crosslink the ADA-GEL. After crosslinking, the CaCl_2_ solution is removed, and the gel is washed three times with PBS.

### Rheological measurements

#### Nonlinear stress-strain relationship

Material properties of hydrogels are measured using a cone-plate shear rheometer (Discovery HR-3, TA Instruments, Milford; geometry: 20 mm diameter, 2° angle, and 54 µm truncation gap) as described previously (34). In brief, collagen and Matrigel samples (85 µl) are loaded and polymerized between the rheometer plates at 37°C for 30 minutes. GelMA (High) and GelMA (Low) samples are photocrosslinked using the formulation-specific UV exposure conditions described above. A solvent trap is used to prevent evaporation during polymerization and measurement. Subsequently, a frequency sweep (0.02-2 Hz) is performed at a strain amplitude of 1%. The stress-strain relationship is then measured by applying a strain ramp from 0% to 100% at a constant strain rate of 1%/s.

#### Cyclic tests

Cyclic tests with repeated loading are conducted using a parallel-plate shear rheometer (Discovery HR-3, TA Instruments, Milford; geometry: 20 mm diameter) equipped with cylindrical porous plastic caps attached to both plates (20 mm diameter, 1 mm height; 3D printed by Shapeways, Livonia) to improve the attachment of the collagen gels and prevent sample slipping. Collagen samples with different concentrations of 0.6 mg/ml, 1.2 mg/ml, or 2.4 mg/ml (550 µl; 1 mm height) are polymerized for 60 minutes between the rheometer plates at 37°C prior to the measurements. For each maximum shear strain amplitude (2%, 4%, 6%, 8%, 10%, or 15%), five consecutive loading cycles are applied at a constant shear rate of 1%/s. The linear stiffness *K* is determined separately for each cycle from the slope of a linear fit to the stress-strain relationship between −1% and +1% strain. The relative stiffness is calculated as *K/K*_0_, where *K*_0_ denotes the stiffness measured during the first cycle at the respective strain amplitude.

### Spheroid traction force measurements

#### Spheroid generation

Traction forces generated from multicellular spheroids in 3D collagen matrices are measured similarly as previously described in (26, 27). Spheroids are cultured in cell-repellent 96-well plates (Greiner Bio-One, Kremsmünster) by typically suspending 4000 cells in 100 µl cell culture medium per well. Within 24 hours, the cells self-assemble into compact spheroids.

#### Spheroid embedding

After 1 day of incubation, the spheroid suspensions from individual wells are pooled in 15 ml Falcon tubes (Corning, New York), with 8-12 spheroids per tube. Excess medium is aspirated, and the spheroids are kept on ice. The collagen solution is prepared on ice and mixed with 4 µm silica beads (5-10 µl per 1 ml of gel; Kisker Biotech, Steinfurt). To position the spheroids within the 3D collagen hydrogels and prevent them from settling onto the bottom of the dish, a two-layer approach is used. First, 1.5 ml of the bead-containing collagen solution is added to each well of a 6-well plate (Corning, New York). Polymerization is initiated by incubating the plate at 37°C with 5% CO_2_. After 15 minutes, 8-12 spheroids from each tube are carefully re-suspended in another 1.5 ml of freshly prepared bead-containing collagen solution and added onto the initial collagen layer. The samples are then polymerized for 1 h at 37°C with 5% CO_2_ to allow complete polymerization of the second collagen layer. Following gelation, 2 ml of pre-warmed cell culture medium is added to each well. A similar two-layer embedding approach is applied to the other hydrogel formulations using the formulation-specific polymerization or crosslinking conditions described above.

#### Timelapse imaging

The samples are transferred to a motorized microscope (Applied Scientific Instrumentation, Eugene) placed inside a tissue culture incubator at 37°C with 5% CO_2_. The equatorial plane of each spheroid is imaged typically every 2 min for 24 h in brightfield mode with a 4x objective (0.13 NA, Olympus, Tokyo) and a CCD camera (Lumenera Infinity 3-6UR, Lumenera Corporation, Ontario).

#### Force reconstruction

From the timelapse images, the cumulative matrix deformations in the collagen gels around each spheroid are measured dynamically using particle image velocimetry (26, 32). Contractile forces are determined from cumulative matrix deformations as previously described in (26). Our approach uses material-specific look-up tables that are derived from finite-element simulations of the experimental setup and account for the nonlinear rheological properties of our collagen hydrogels measured by shear rheology and extensional strain experiments in (34). Using these tables, cell-generated forces are reconstructed from the amplitude of local matrix deformations and their respective distances from the spheroid, both normalized to spheroid size. The force values obtained from all local matrix deformations with a minimum distance of 2 spheroid radii from the spheroid center are then averaged per spheroid and per time step. For the analysis of neighboring spheroids (Figure 6), the force values are calculated from the matrix deformations in a 5° angular cone perpendicular to the line connecting the spheroids.

### Detection of force oscillations

To detect minute-scale force oscillations during the 24 h measurement period, a two-step analysis is performed. First, a Butterworth high-pass filter (48) with a cutoff frequency of 1/(60 min) is applied to the force time curves of each spheroid to remove slow baseline changes and retain force fluctuations with periods shorter than 60 min. Second, force pulses are detected as local minima in the high-pass-filtered force curves using the Python package SciPy (64). Successive minima are required to be separated by at least 5 min and to have an absolute amplitude of at least 3 µN. The pulse frequency of each spheroid is calculated by dividing the number of detected pulses by the analyzed measurement duration.

### Drug screening measurements

For drug screening experiments, pharmacological activators and inhibitors are added during spheroid traction force measurements using two experimental protocols. First, in the dynamic protocol, drugs are added to the culture medium at time points between 9 and 15 h during the 24 h measurement period. The overall contractile force, pulse frequency, and pulse amplitude are quantified within 5 h windows before and after compound addition. Second, in the static protocol, compounds are added to the culture medium at the beginning of the 24 h measurement period, after the gels have polymerized for 1 h. The overall contractile force, pulse frequency, pulse amplitude, and matrix invasion are then quantified over the entire 24 h measurement period. The corresponding vehicle controls contain the same final concentration of solvent (PBS, DMSO, or H_2_O) as the compound-treated samples. SI Fig. 9 shows an overview of the used compounds and treatment conditions.

### Matrix invasion measurements

After 24 h in collagen, spheroids are fixed, permeabilized, and stained for F-actin and nuclei as follows. Cell culture medium is aspirated, and the collagen gels are washed twice with PBS. A total of 1 ml of fixative solution containing 0.2% paraformaldehyde in PBS (Thermo Fisher, Waltham) and 2% glutaraldehyde in sodium phosphate buffer (Merck, Darmstadt) is added to each well for 20 min at room temperature. The fixative is removed, and the collagen gels are washed twice with PBS. The samples are subsequently permeabilized with 0.2% Triton X-100 (Carl Roth, Karlsruhe) for 1 h at room temperature. The permeabilization solution is then removed, and the samples are washed twice with PBS. Next, 1 ml of 0.05% Triton X-100 is added for 1 h at room temperature. After washing twice with PBS, the samples are incubated with 500 µl of 0.05% Triton X-100 containing Hoechst dye (2.5 µg/ml, Sigma-Aldrich, St. Louis) and TRITC-conjugated phalloidin (20 µg/ml, Sigma-Aldrich, St. Louis) for 1 h at room temperature protected from light. Samples are washed twice with PBS, after which 2 ml of PBS is added. The equatorial plane of each spheroid is imaged using a fluorescence microscope (Leica DMI600 CS, Leica, Wetzlar) equipped with a 5x objective (Leica, Wetzlar) and a CCD camera (Lumenera Infinity 3-6UR, Lumenera Corporation, Ontario). To quantify differences in matrix invasion among drug treatments, the actin-positive cell area is segmented using a fixed intensity threshold applied identically to images of drug-treated and control spheroids. The relative change of segmented area compared to control spheroids is used as a measure of matrix invasion and compared between treatment groups. Images of the collagen fiber structure are acquired using confocal reflection imaging with an upright laser-scanning confocal microscope (Leica SP5, Leica, Wetzlar) equipped with a 20x dip-in objective (HCX APO L 20x/1.00 W, Leica, Wetzlar).

### Microtissue pillar assay

Microtissues from primary human breast cancer-associated fibroblasts are generated by seeding a cell-collagen suspension into wells containing two flexible PDMS pillars, around which the microtissues self-assemble, as described previously in (28, 29). Briefly, custom PDMS pillar arrays (curing agent-to-base ratio of 1:26; Sylgard184, Sigma-Aldrich, St. Louis) are filled with a bottom layer of collagen (0.6 mg/ml collagen; 6 µl per well) and polymerized for 1 hour at 37°C and 5% CO_2_. A second layer consisting of 6 µl of collagen solution (0.6 mg/ml) containing 4000 cells per well is added and polymerized for another 1 hour under the same conditions. Finally, 1 ml of pre-warmed cell culture medium is added to the pillar array. Images of the equatorial plane of the tissue and the top of the PDMS pillars are acquired every 2 min for 24 h in bright-field mode using a 4x objective (0.13 NA, Olympus, Tokyo) and a CCD camera (Lumenera Infinity 3-6UR, Lumenera Corporation, Ontario) in a microscope (Applied Scientific Instrumentation, Eugene) placed inside a tissue culture incubator (37°C and 5% CO_2_). The pillar deflection at the top of each pillar is tracked using the Python package ClickPoints (65). The pillar deflection at the tissue plane is then calculated from the deflection measured at the top of the pillar, the tissue height, and Euler–Bernoulli beam theory (28, 29). This deflection, together with the effective spring constant of the pillar, is used to calculate the contractile force generated by each microtissue (28, 29).

Microtissues from patient-derived human rectal adenocarcinoma-associated fibroblasts are generated as described previously (30, 66). In this system, microtissues are engineered within polydimethylsiloxane (PDMS; Sylgard 184, Dow Corning) microwells containing two T-shaped cantilevers. Microwells are obtained by PDMS replication of SU-8-based masters that are microfabricated as previously described (66–68). The cantilever spring constant k is calibrated using a capacitive MEMS force sensor mounted on a micromanipulator, as described previously (30, 67–69). The resulting spring constant is k = 0.45 *±*0.10 N/m. PDMS microwells are sterilized in 70% ethanol and treated with 0.2% (w/v) Pluronic F127 (Sigma) for 2 min to reduce cell adhesion. A cooled suspension containing 300.000 CAF cells in 1 ml of neutralized liquid rat-tail collagen I (2 mg/ml; Advanced Biomatrix, Carlsbad) is then added to the microwells. The array is centrifuged to sediment the cells into the microwells. The constructs are polymerized at 37°C for 9 min, followed by the addition of culture medium. Microtissues are maintained in the incubator for 24 h prior to the experiments. During this period, the cells spread and compact the collagen matrix to form microtissues suspended between the tips of each pair of cantilevers. The force generated by each microtissue is determined from cantilever deflection as described previously (66–68). Brightfield images are acquired at intervals of 5 to 30 s, and the positions of the cantilever tips are tracked using a previously developed MATLAB script (68). Tracking results are checked visually, and incorrectly tracked positions are manually corrected or excluded from the analysis if reliable correction is not possible. The force F generated by the microtissue is calculated from the average deflection dAVG of the two cantilevers and the spring constant k as follows: F = k *·*dAVG. Only tissues that remain uniformly anchored to the tips of both cantilevers throughout the experiment are included in the analysis.

## Data Availability

The traction force microscopy software and the collagen-specific look-up tables for force reconstruction are available on GitHub as an open-source Python package (26, 70). Figures are created using the Python package Pylustrator (71) and MetBrewer (72). The data from this study are available upon reasonable request from the corresponding author.

## Author contributions

BF, TB and IT supervised the study. RS and PS established the primary cultures of patient-derived breast cancer-associated fibroblasts. CM, RG and DB developed the spheroid traction force analysis method. DB and LV conducted spheroid traction force experiments. DMV, MB, and GC conceived and designed the microtissue experiments using adenocarcinoma-associated fibroblasts, and OZ performed these experiments. DB and DK performed micro-tissue experiments using breast cancer-associated fibroblasts. AH, SB and DB conducted the rheological measurements. ARB and JR developed and prepared the alginate-based hydrogels. LS, GW, HH, WHG, DB, LF, and LW established and conducted experiments using mouse skin fibroblasts. TB, LV and DB analyzed the data, created figures and videos. BF, DB and TB wrote the manuscript with feedback and contributions from all authors.

## ACKNOWLEDGEMENTS

This work was funded by grants from the German Science Foundation (DFG; project 326998133 TRR-SFB 225 subprojects A01, B09, C02, and C05; project 460333672 – CRC 1540 – subprojects A01 and C05) and the National Institutes of Health (HL120839). T.B. and B.F. acknowledge funding from the ANR/DFG CONTRACTILE project, grant ANR-23-CE13-0037 of the French Agence Nationale de la Recherche (ANR). We thank Johanna Ivaska (University of Turku, Finland) and John Eriksson (Åbo Akademi University, Turku, Finland) for kindly providing the vimentin wild-type and vimentin-deficient mouse embryonic fibroblasts. We also thank Ricardo Henriques Laboratory for the use of the LaTeX layout under Creative Commons 4.0 License.

---

**Extended Data Figure 1:**
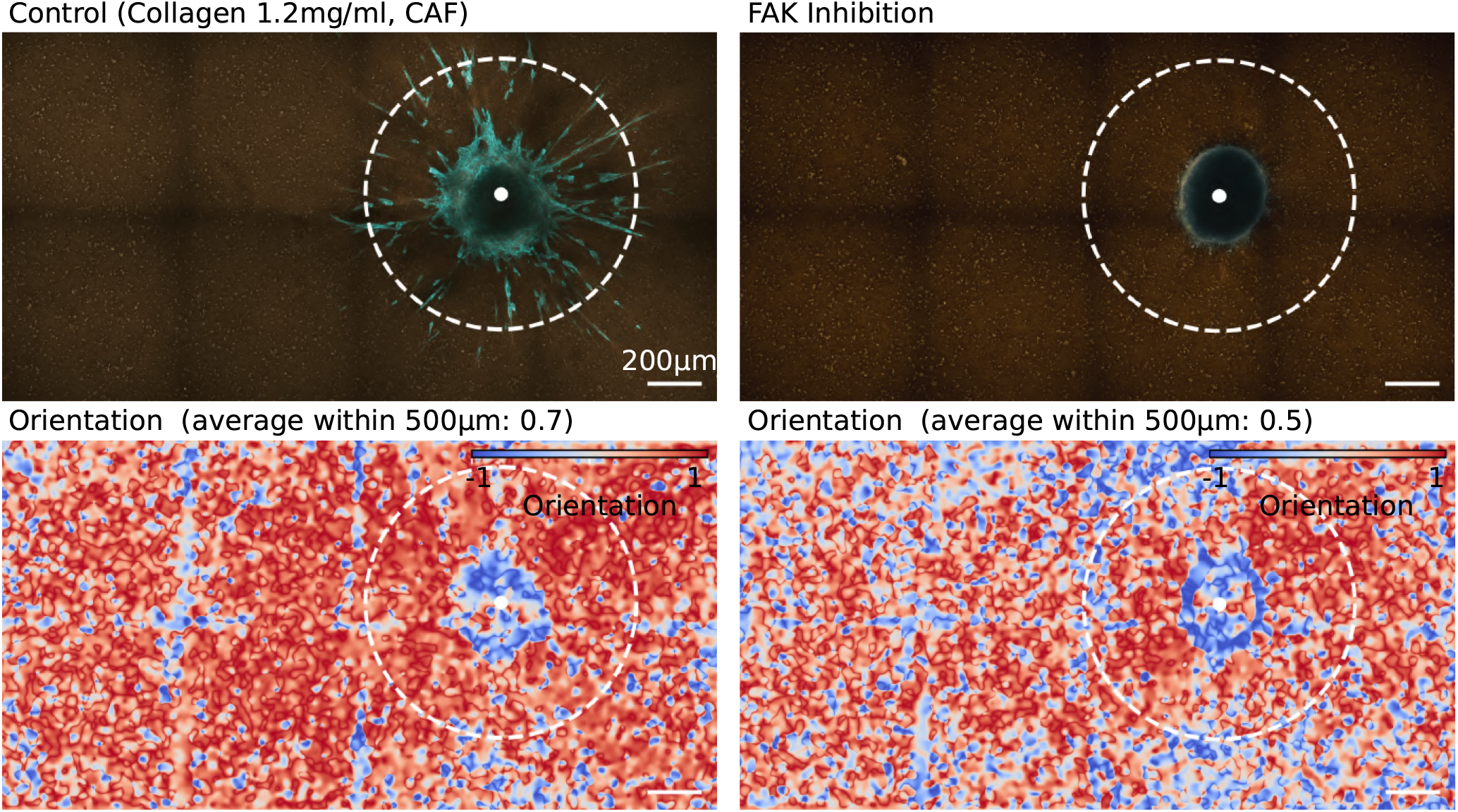
Cell invasion and collagen alignment around CAF spheroids with and without FAK inhibition. Spheroids generated from 4000 primary breast cancer-associated fibroblasts are shown after 24 h in 1.2 mg/ml 3D collagen matrices. The left panel shows a control spheroid, whereas the right panel shows a spheroid treated with 3 µM defactinib, an inhibitor of focal adhesion kinase (FAK), during the measurement period. Compared with the control, the defactinib-treated spheroid shows reduced collagen fiber alignment and little to no cell invasion. F-actin is stained with TRITC-conjugated phalloidin and displayed in blue. Collagen fibers are imaged by confocal reflection microscopy and displayed in brown. The images are stitched from eight adjacent fields of view. The lower panels show collagen fiber orientation relative to the radial direction towards the spheroid center, indicated by the white dot. Fiber orientation is quantified by structure tensor analysis as described previously (31). Here, a value of 1 indicates radial alignment towards the spheroid center, whereas a value of −1 indicates alignment perpendicular to the radial direction. Orientation is calculated using a window size of 10 µm from a maximum-intensity projection of a 10 µm-thick confocal reflection stack centered on the spheroid equatorial plane. Mean orientation values within a radial distance of 500 µm from the spheroid center (dotted line) are shown above the images.

**Extended Data Figure 2:**
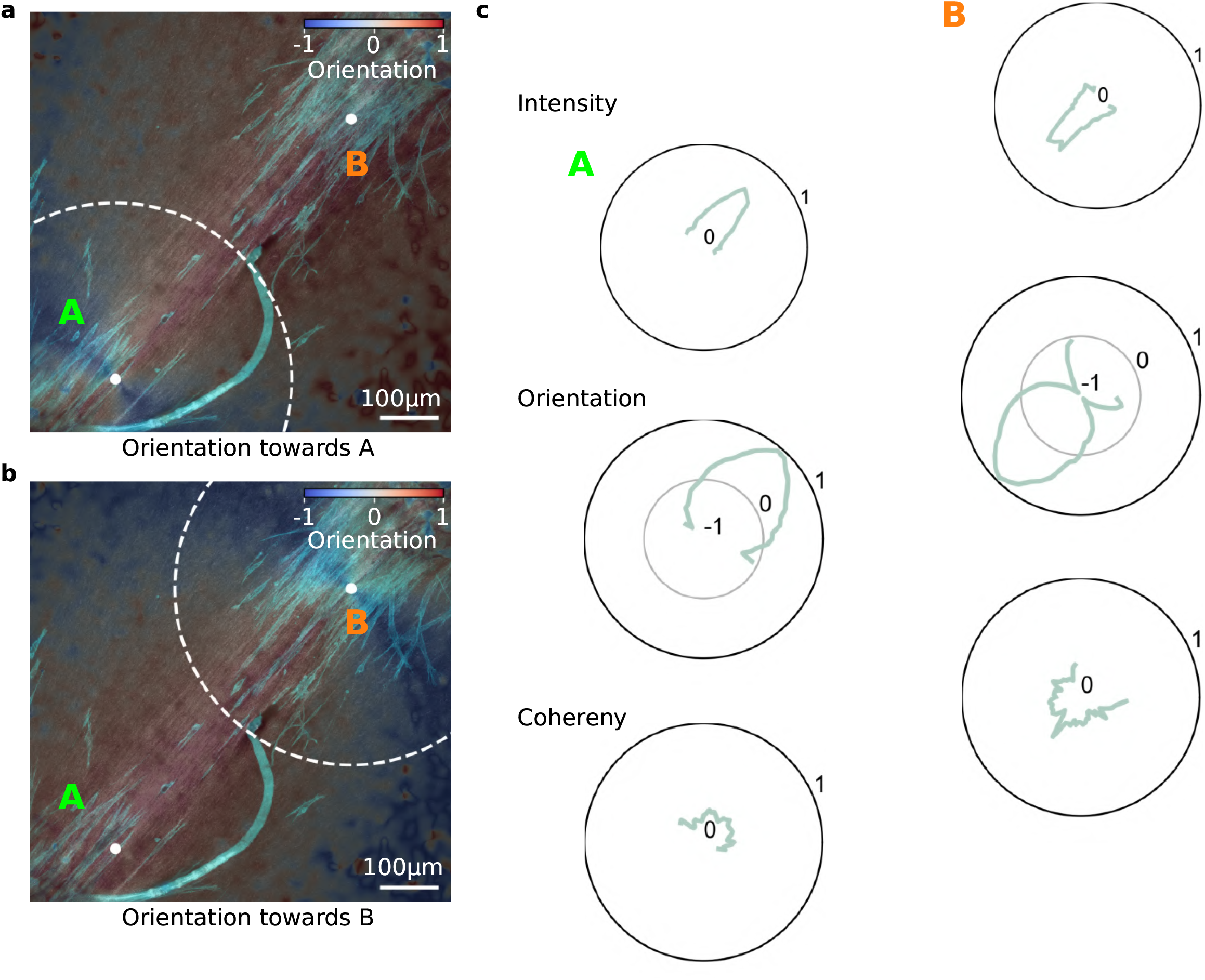
Mechanical interaction between adjacent CAF spheroids. Two spheroids, each generated from 4000 breast cancer-associated fibroblasts (CAFs), are embedded in 1.2 mg/ml collagen gels for 24 h. **a**,**b**, Maximum-intensity projections of confocal reflection images showing collagen fibers within an 8 µm-thick z-stack around the spheroid equatorial plane are overlaid with maximum-intensity projections of confocal fluorescence images showing the actin-stained cells within a 302 µm-thick z-stack. F-actin is stained using 20 µg/ml TRITC-conjugated phalloidin (Sigma-Aldrich, St. Louis). The blue-to-red color scale indicates collagen fiber orientation relative to spheroid A (**a**) and spheroid B (**b**). A value of 1 indicates radial alignment towards the respective spheroid center, whereas a value of −1 indicates alignment perpendicular to the radial direction. Contractile remodeling of the collagen matrix generates an aligned collagen fiber band connecting the two spheroids. **c**, Polar plots show collagen fiber intensity, orientation, and coherence around spheroid A (left) and spheroid B (right), averaged within a radial distance of 300 µm from the respective spheroid center (dotted lines in **a**,**b**). Figure reproduced from SI Fig. 12 in (31) under CC BY 4.0.

**Extended Data Figure 3:**
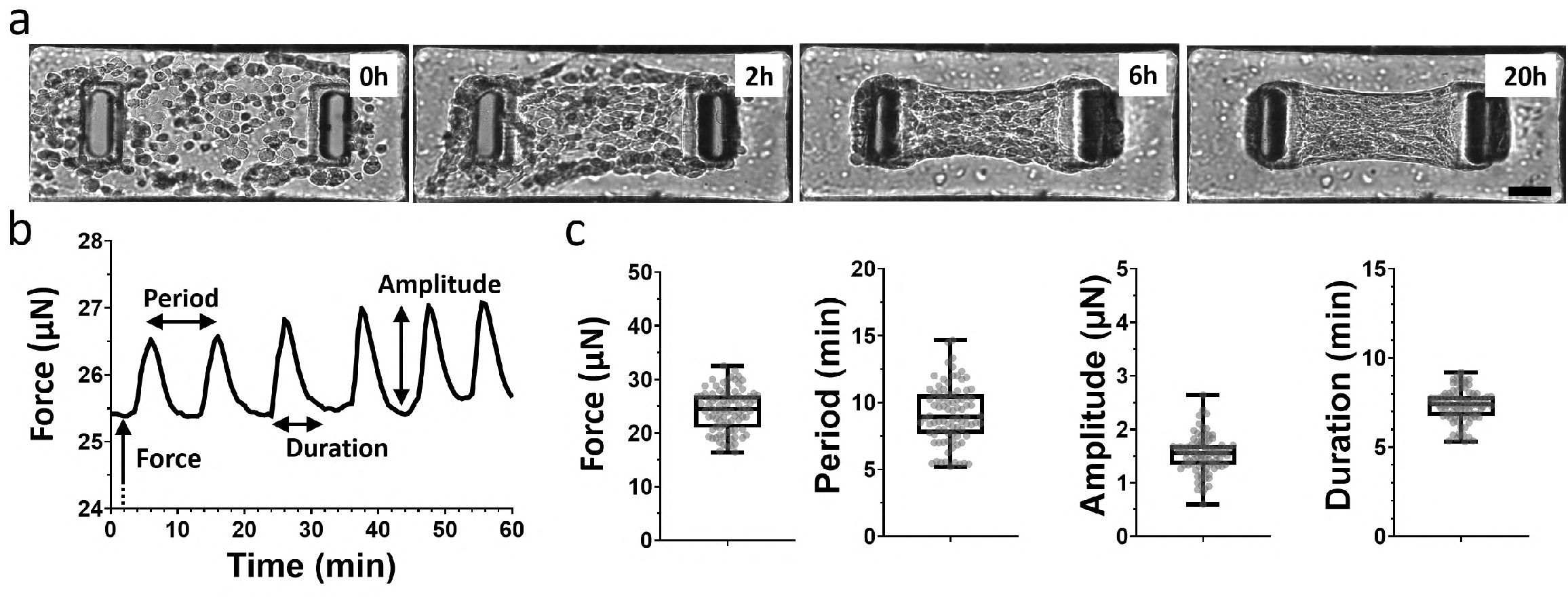
Force oscillations in microtissues from human adenocarcinoma fibroblasts. **a**, Brightfield images showing the self-assembly of microtissues composed of 300 patient-derived human cancer-associated fibroblasts (CAFs), isolated from a Lieberkühnian adenocarcinoma of the upper rectum, embedded in 2 mg/ml collagen I over 20 hours. Scalebar indicates 100 µm. **b**, Representative force trace from a microtissue after 24 h in culture, showing repeated force oscillations similar to those observed in other human CAF microtissues and in the spheroid system (Figure 1). **c**, Force, oscillation period, oscillation amplitude, and the pulse duration are averaged for 87 different microtissues. For each box plot, the box extends from the 25th to 75th percentiles, the median is plotted as a line inside the box, the whiskers extend to the minimum and maximum values, and the individual data points are overlaid.

**Extended Data Figure 4:**
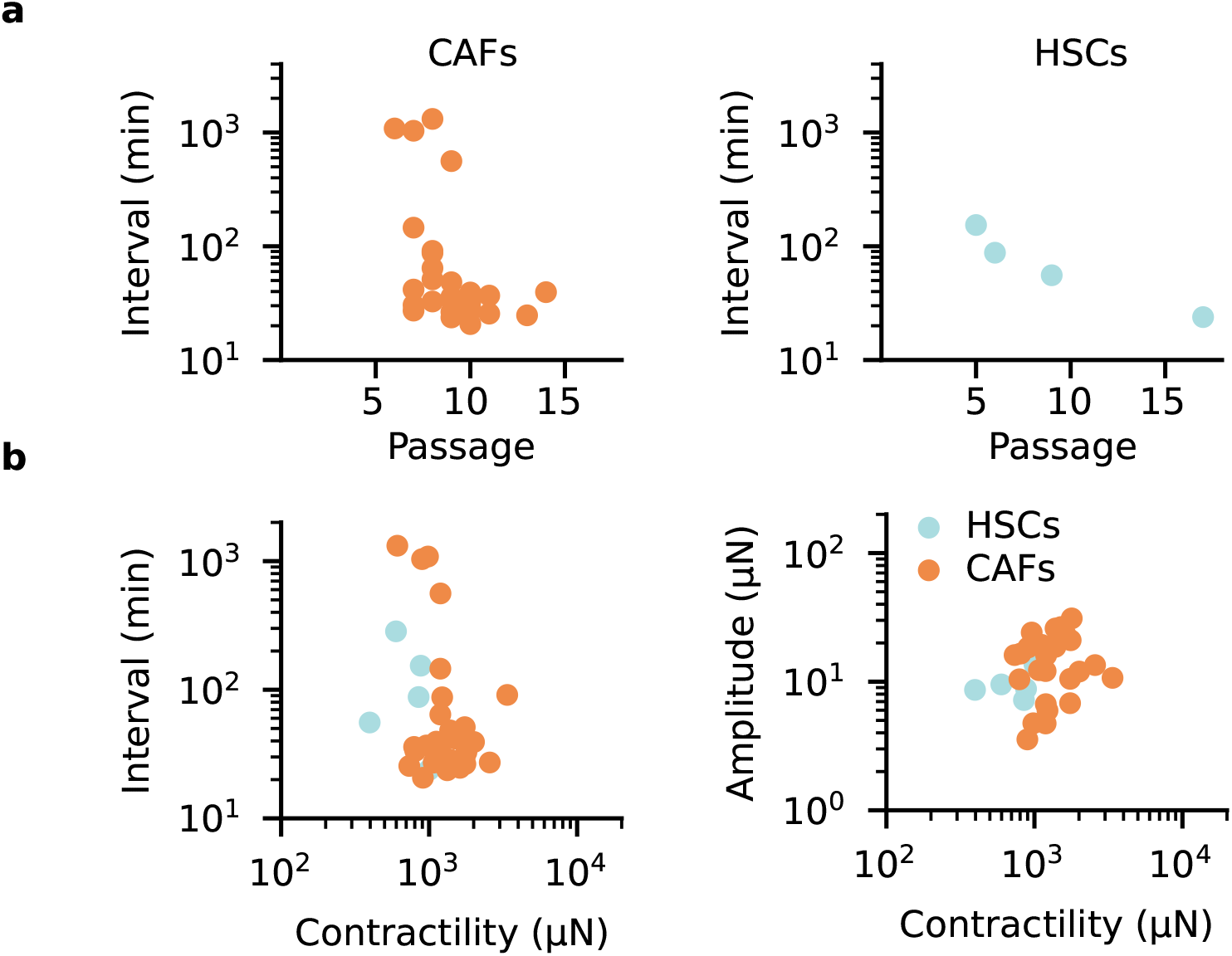
Spheroid force oscillations scale with contractility and cell culture passage. Time interval between force oscillations in breast cancer-associated fibroblast (CAF) and hepatic stellate cell (HSC) spheroids at different cell passage numbers. Each point represents the mean across 4–10 spheroids from an individual measurement. At higher passage numbers, both CAF and HSC spheroids show shorter intervals between force oscillations. **b**, Time interval between force oscillations (left) and_2_ force-oscillation amplitude (right) as a function of spheroid contractility. Each point represents the mean across 4–10 spheroids from an individual measurement.

**Extended Data Figure 5:**
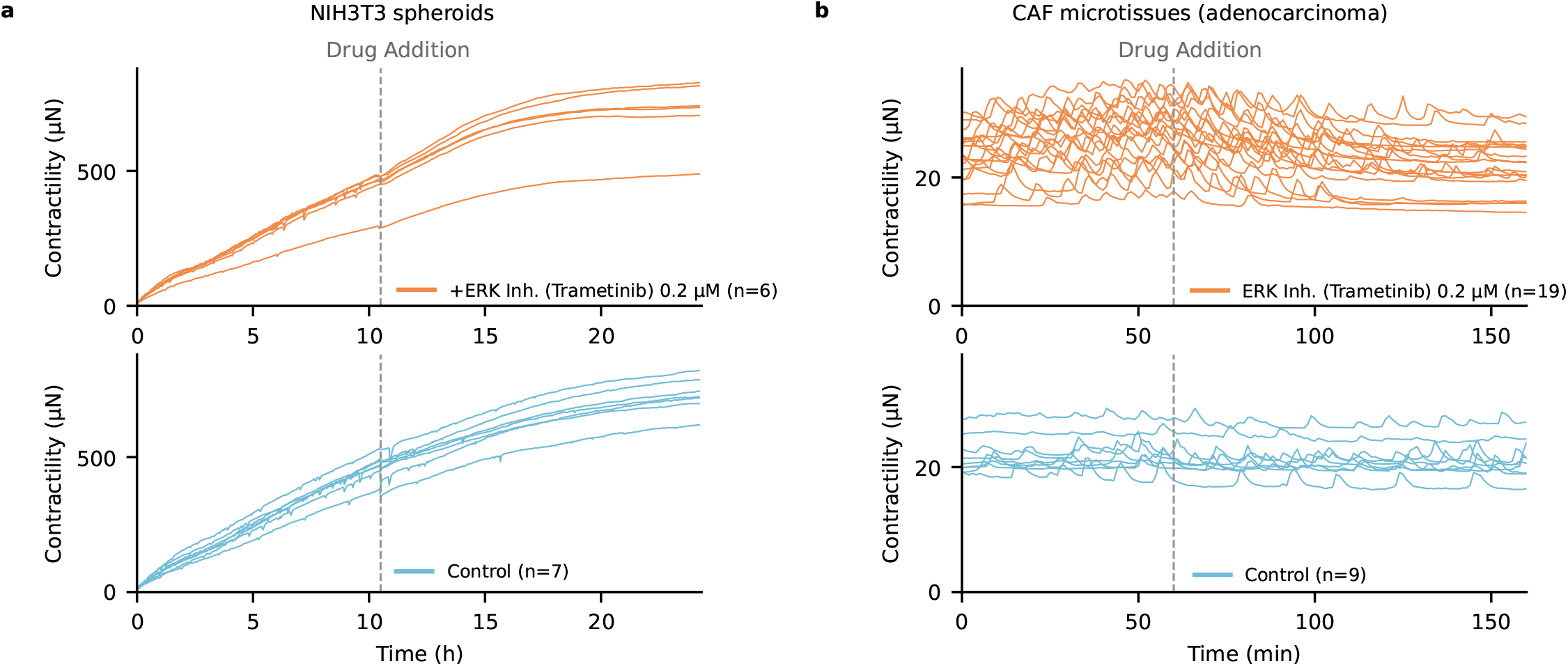
MEK/ERK pathway inhibition in NIH3T3 spheroids and human CAF microtissues a, Traction force measurements of spheroids generated from 4000 NIH3T3 cells. Trametinib, a MEK/ERK inhibitor, (0.2 µM; top) or the corresponding vehicle control (bottom) is added during the measurement at the time indicated by the gray dashed line. N indicates the number of spheroids. **b**, Traction force measurements of microtissues generated from 300 patient-derived human rectal adenocarcinoma-associated fibroblasts. As in **a**, 0.2 µM Trametinib (top) or the corresponding vehicle control (bottom) is added during the measurement at the time indicated by the gray dashed line. N indicates the number of microtissues.

**Extended Data Figure 6:**
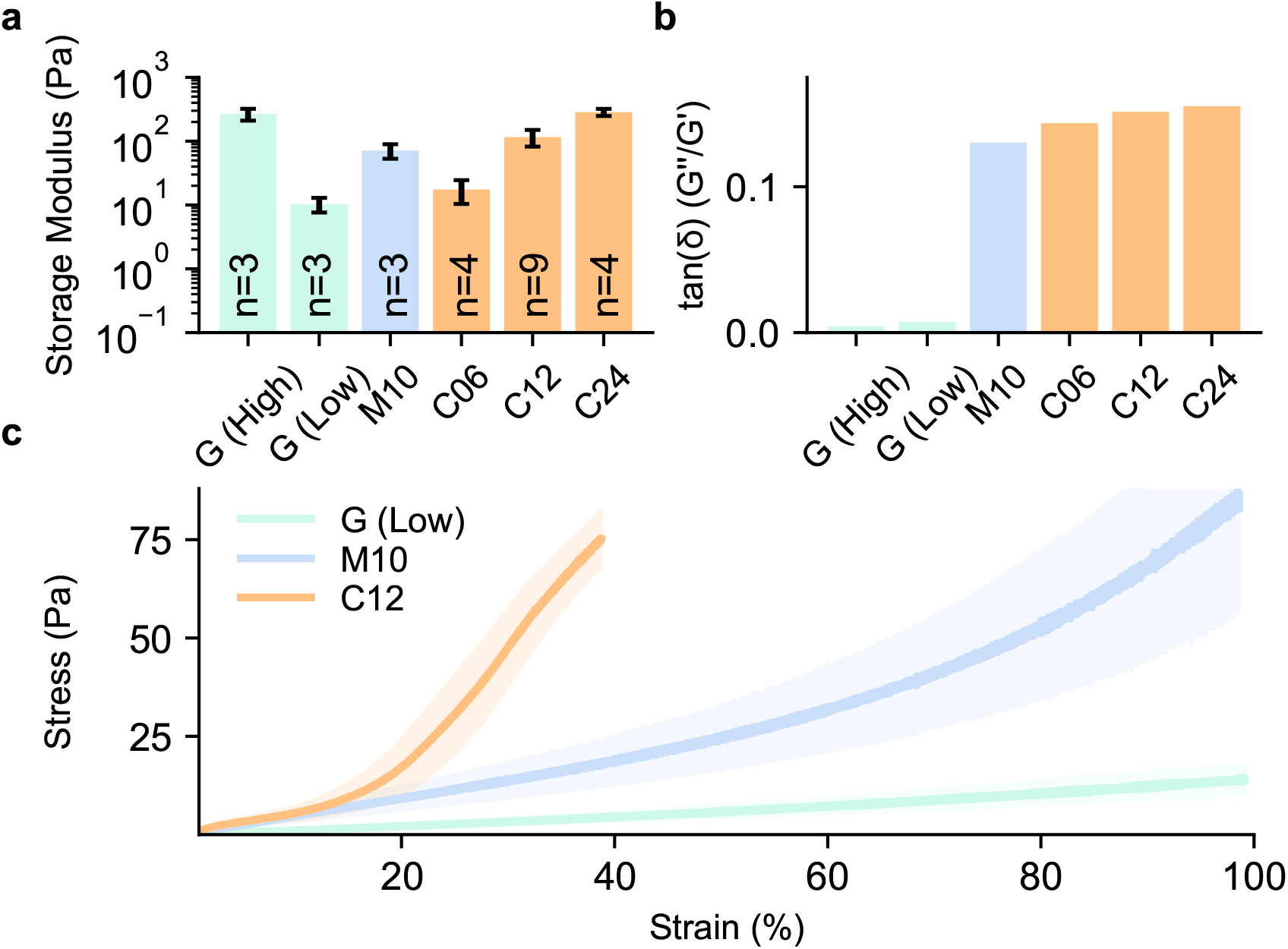
Rheological properties of matrices with different degrees of strain stiffening and viscoelasticity. Shear rheological measurements of different matrices with various degrees of strain stiffening and viscoelasticity. GelMA is tested using two formulations, GelMA (High) and GelMA (Low) (see Methods). Matrigel is measured at a concentration of 10 mg/ml (M10), whereas collagen type I gels are measured at concentrations of 0.6 mg/ml, 1.2 mg/ml, and 2.4 mg/ml (C06, C12, and C24). **a**, Storage modulus measured at 0.02 Hz and 1% strain amplitude. Bars indicate the mean*±* sd. **b**, The loss tangent, calculated as the ratio of the loss modulus (G”) to the storage modulus (G’), quantifies matrix viscoelasticity at 2 Hz and 1% strain amplitude. Matrigel and collagen show viscoelastic behavior dominated by an elastic response, whereas GelMA exhibits an almost purely elastic response with a negligible viscous contribution. **c**, Shear stress response to a continuous strain ramp (1%/s). Collagen exhibits pronounced nonlinear strain stiffening beginning at *≈* 10% strain, whereas Matrigel shows less pronounced strain stiffening beginning at *≈* 50% strain. GelMA does not show nonlinear strain-stiffening.

**Extended Data Figure 7:**
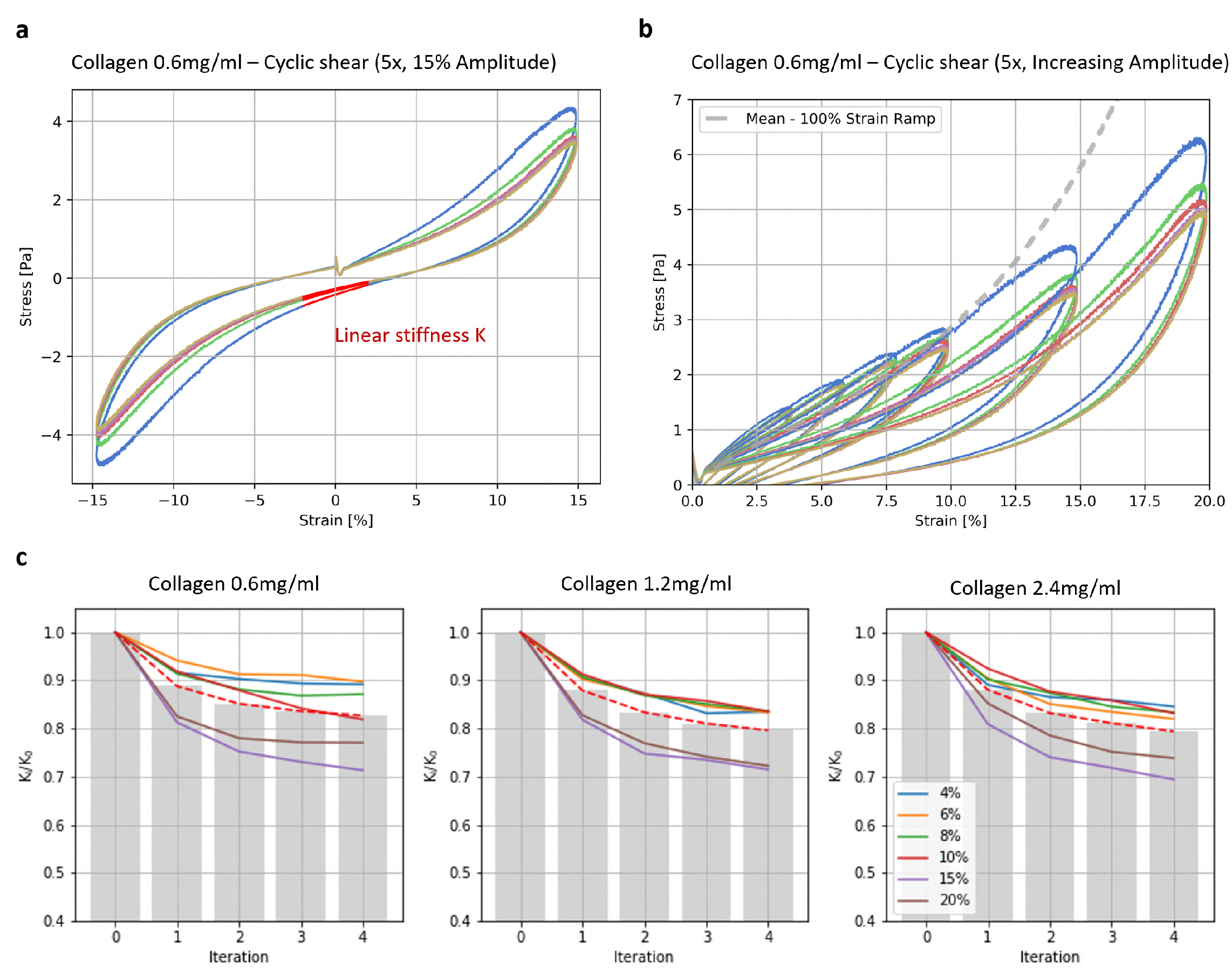
Uniform cyclic weakening of collagen gels at different concentrations. Collagen samples with concentrations of 0.6 mg/ml, 1.2 mg/ml, and 2.4 mg/ml are polymerized between the plates of the shear rheometer. The shear stress-strain relationship is then measured over five consecutive loading cycles (**a**) at increasing maximum strain amplitudes (2%, 4%, 6%, 8%, 10%, and 15%; see **b**). For each amplitude and cycle, the linear stiffness is determined from the slope of a linear fit over the strain range from −1% to +1%. The gray dotted line in **b** shows the mean stress-strain response of collagen gels without cyclic loading (continuous 100 % strain ramp) and is comparable to the initial loading cycles at low strain amplitudes. **c**, Relative collagen stiffness is calculated by dividing the linear stiffness of each cycle, *K*, by the stiffness of the first cycle, *K*_0_ (K/K_0_). At strain amplitudes above 2%, relative collagen stiffness progressively decreases with increasing cycle number. The degree of weakening increases with strain amplitude, reaching a stiffness decrease of approximately 25-30% at the highest applied strain amplitude. Red dotted lines indicate the average decrease in relative stiffness across the displayed strain amplitudes. No weakening is observed at a strain amplitude of 2%. The magnitude of cyclic weakening is similar across the different collagen concentrations.

**Extended Data Figure 8:**
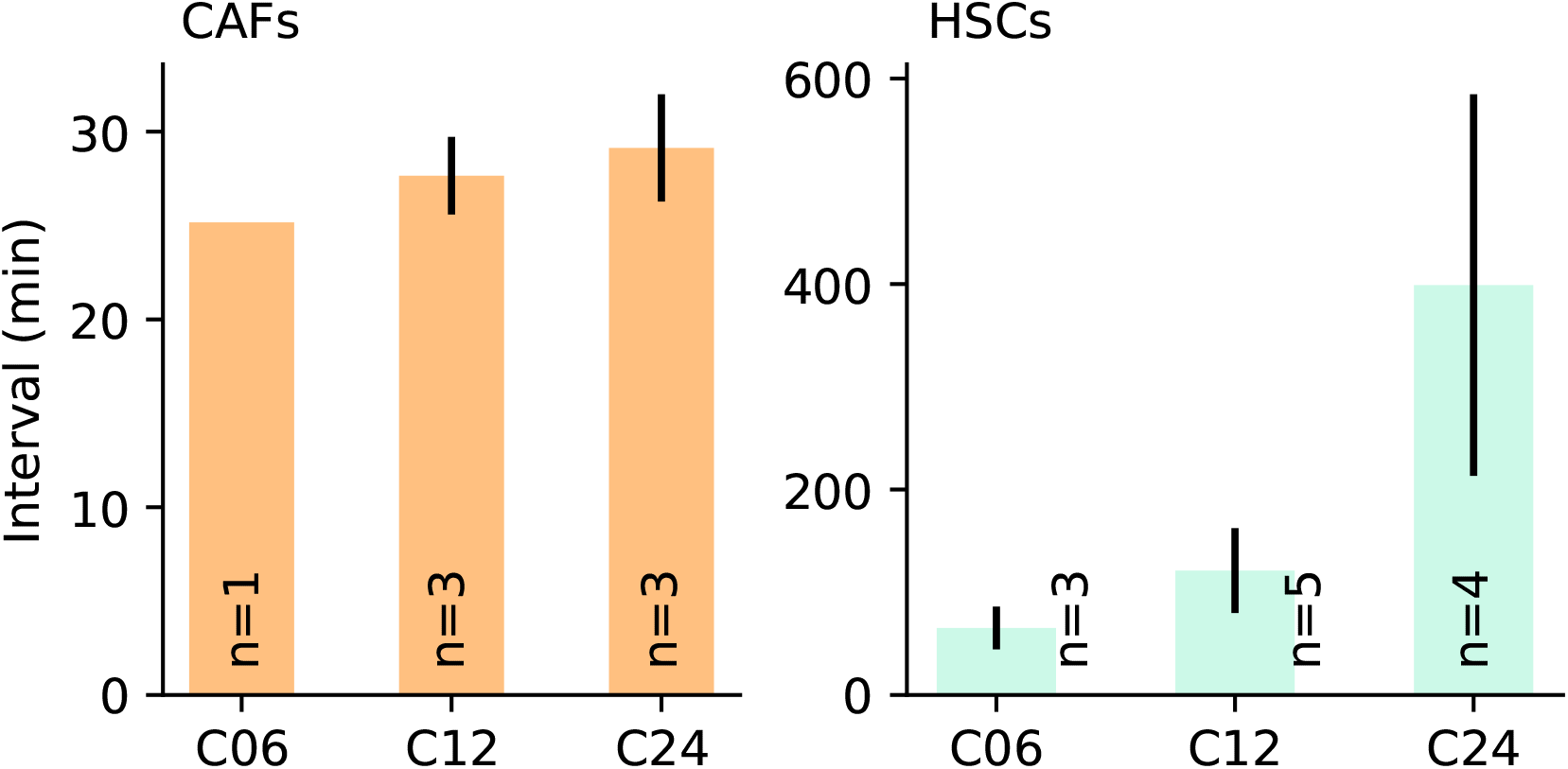
Spheroid force oscillation for different collagen concentrations. Time interval between force oscillations in breast cancer-associated fibroblast (CAF) spheroids (left) and hepatic stellate cell (HSC) spheroids (right), measured over 24 h in collagen. Bars indicate the mean *±*se of individual experiments, each containing multiple spheroids. The interval between force oscillations increases slightly with increasing collagen concentration, corresponding to a decrease in oscillation frequency. C06, C12, and C24 refer to collagen gels with concentrations of 0.6 mg/ml, 1.2 mg/ml, and 2.4 mg/ml, corresponding to storage moduli of 14 Pa, 84 Pa, and 200 Pa, respectively. Storage moduli are determined by shear-rheological measurements at 0.02 Hz and 1% strain as described previously in (31) (see Methods and SI Fig.6).

**Extended Data Figure 9:**
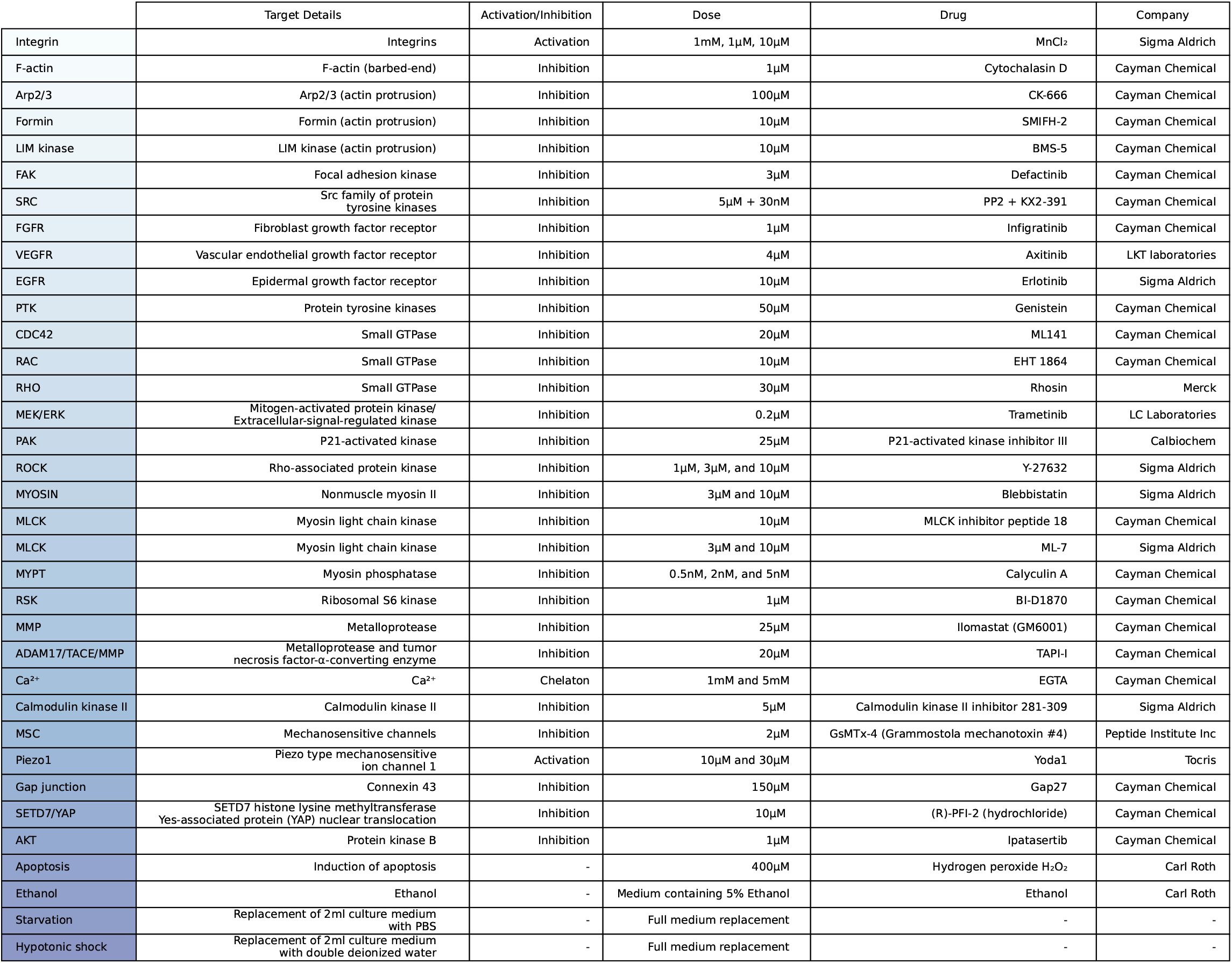
Table of compounds used for drug screening experiments. Overview of the pharmacological compounds used in the spheroid traction force experiments. The table lists the compound abbreviations, molecular targets, working concentrations, compound names, suppliers, and classification as activators or inhibitors.

**Extended Data Figure 10:**
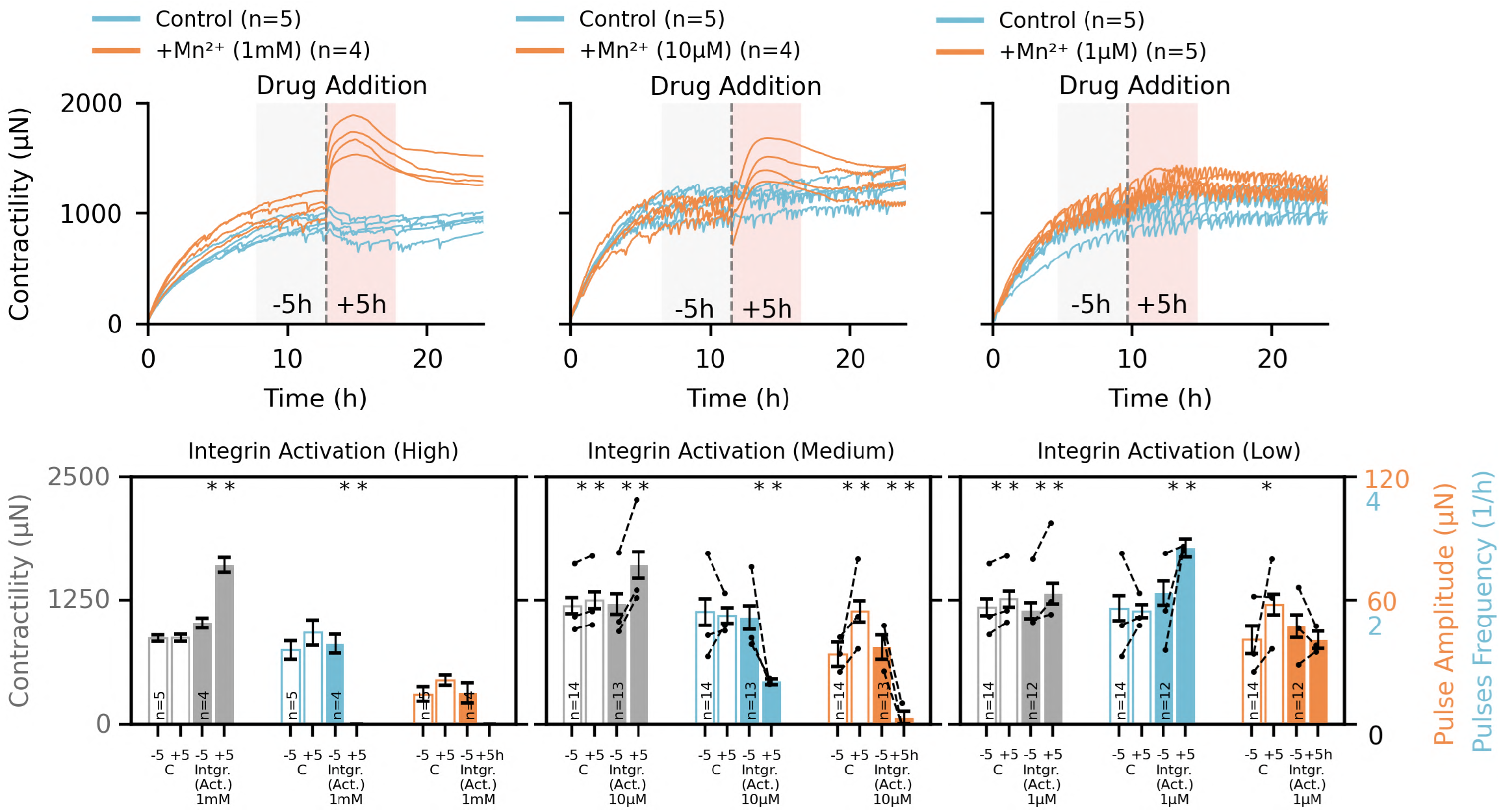
Effects of integrin activation and FAK inhibition on force oscillations. Top,Mn^2+^ is used to activate integrins, whereas focal adhesion kinase (FAK) is inhibited to disturb focal adhesion dynamics during spheroid traction force measurements. Contractile forces are shown following treatment with different concentrations of Mn^2+^ (orange) and for the corresponding control spheroids (blue). The gray dotted line indicates the time of compound addition, and n indicates the number of spheroids. At higher Mn^2+^ concentrations, spheroids show a transient peak-like increase in force (left), whereas lower concentrations increase the frequency of force oscillations (right). **Bottom**, Contractility (gray), pulse frequency (blue), and pulse amplitude (orange) are calculated for each experiment by averaging across all spheroids within 5 h windows before and after drug addition (gray and orange shaded areas, respectively). Bars show the mean *±*se across experiments, and black dots indicate the means of individual experiments. Filled bars represent experiments with compound addition, whereas unfilled bars represent the corresponding vehicle controls containing the same final solvent concentration. The compound names and their molecular targets are indicated below and above the bars, respectively (seeFigure 2and SI Fig.9 for additional details). Significance is determined using a paired t-test, where * indicates p*<*0.05 and ** indicates p*<*0.01.

**Extended Data Figure 11:**
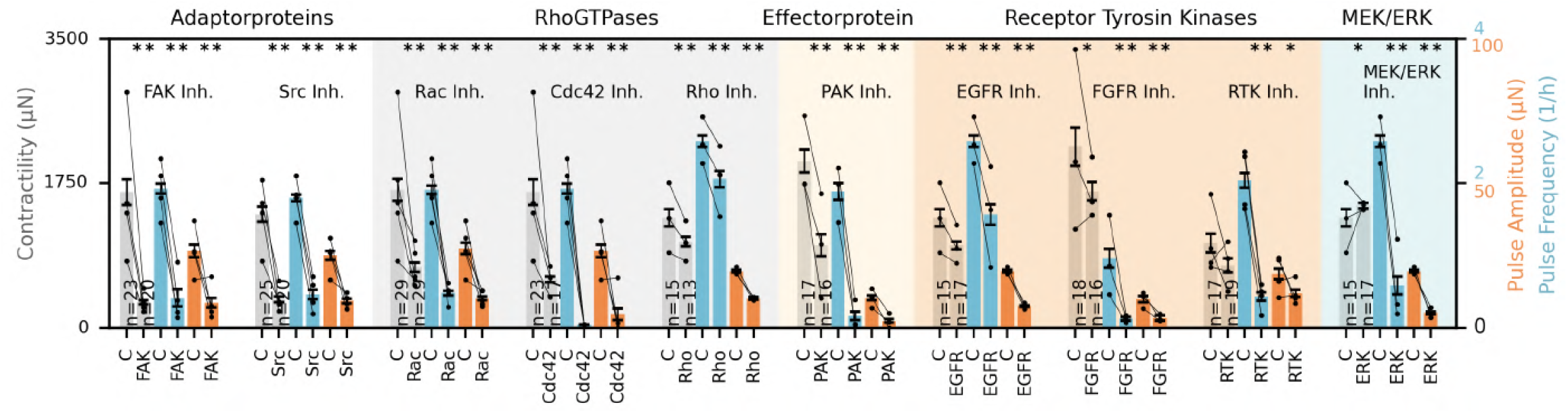
Effects of ERK pathway-associated signaling inhibitors on spheroid contractility and force oscillations. To investigate the signaling network regulating force oscillations, pharmacological inhibitors are used to target components within the ERK pathway throughout the full 24 h spheroid traction force measurements (seeFigure 3). Spheroids are treated with different compounds targeting focal adhesion kinase (FAK), Src family kinases (Src), Rho GTPases (Rac, Cdc42, and Rho), p21-activated kinase (PAK), fibroblast growth factor receptor (FGFR), and MEK/ERK signaling (see SI Fig.9for additional details). Spheroid contractility (gray), pulse frequency (blue), and pulse amplitude (orange) are quantified for drug-treated and corresponding control (C) spheroids over the 24 h measurement period. Bars show the mean*±* se across at least three individual experiments. For each experiment, values are first averaged across all measured spheroids. The total number of spheroids is indicated by n, whereas black dots represent the means of individual experiments. Lines connect corresponding drug-treated and control experiments. The respective compound and molecular target are indicated above the bars. Significance is determined using a paired t-test, where * indicates p*<*0.05 and ** indicates p*<*0.01.

**Extended Data Figure 12:**
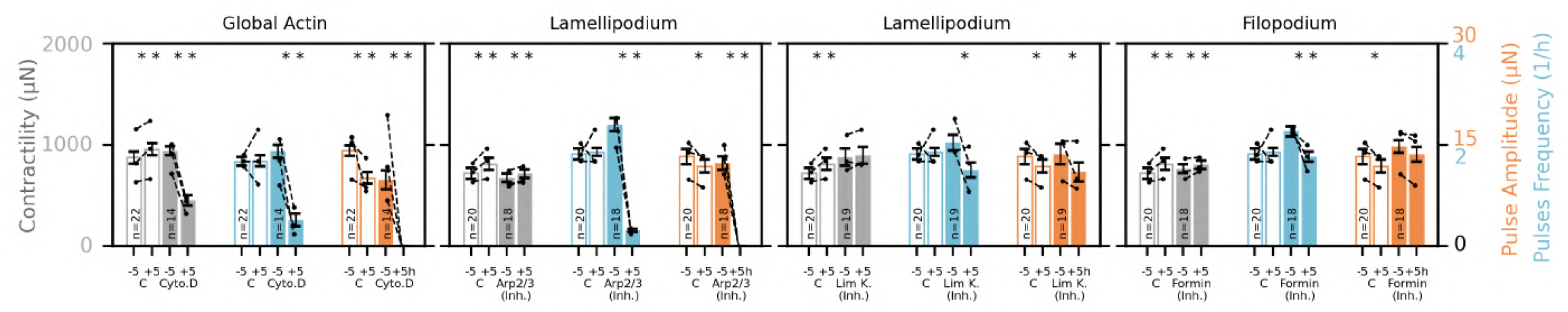
Effects of actin cytoskeleton inhibitors on contractility and force oscillations. During spheroid traction force measurements, cytochalasin D is used to broadly disrupt actin polymerization, whereas a formin inhibitor is used to perturb linear actin assembly associated with filopodia. Inhibitors of the Arp2/3 complex and LIM kinase (LIMK) are used to perturb branched actin assembly and cofilin-dependent actin turnover, respectively, both of which contribute to lamellipodial dynamics (seeFigure 2and SI Fig.9for additional details). Contractility (gray), pulse frequency (blue), and pulse amplitude (orange) are calculated for each experiment by averaging across all spheroids within 5 h windows immediately before and after drug addition. Bars show the mean *±*se across experiments, and black dots indicate the means of individual experiments. Filled bars represent experiments with drug addition, whereas unfilled bars represent the corresponding vehicle controls containing the same final solvent concentration. Compound names and their molecular targets are indicated below and above the bars, respectively. Significance is determined using a paired t-test, where * indicates p*<*0.05 and ** indicates p*<*0.01.

**Extended Data Figure 13:**
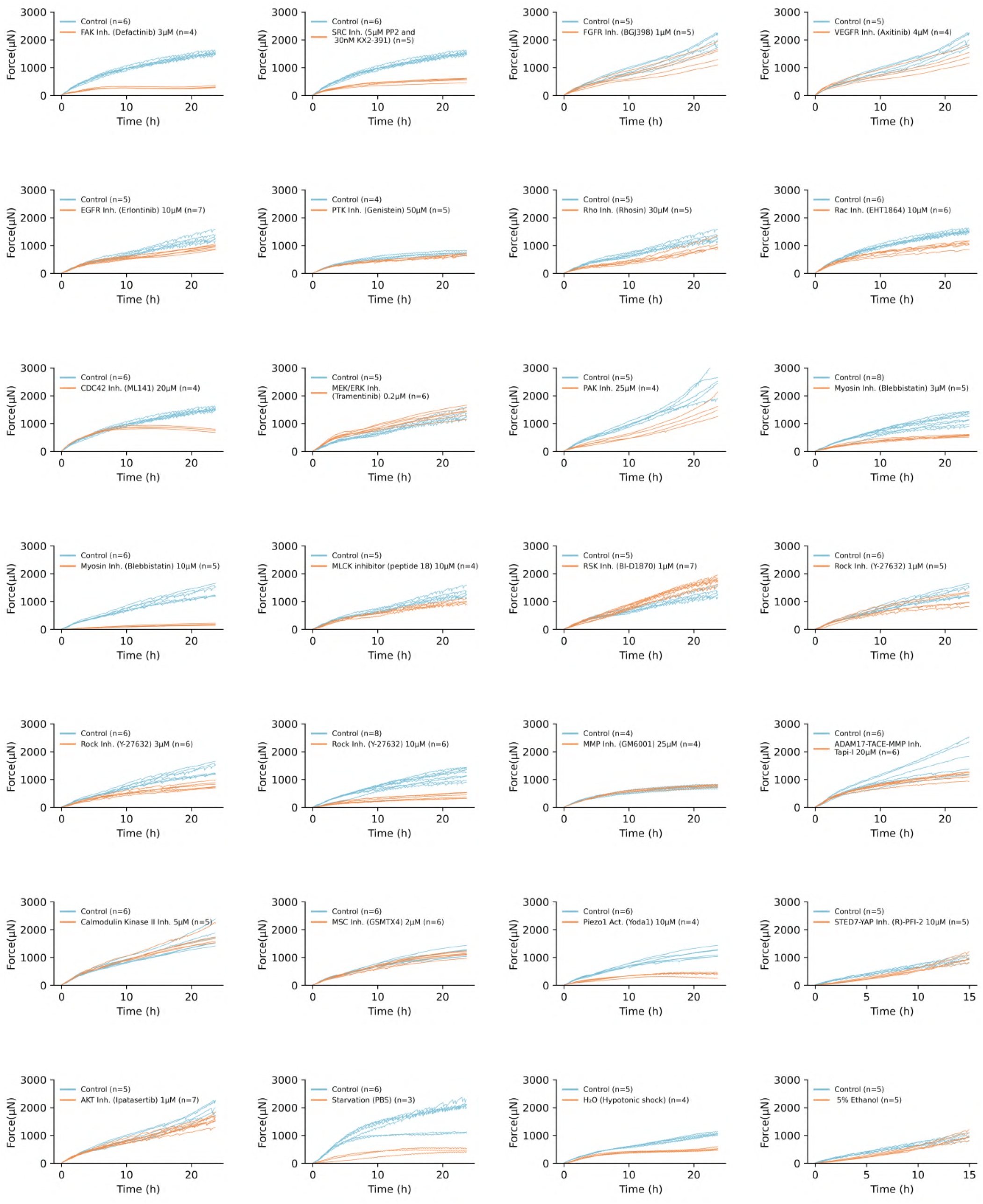
Spheroid force development under static drug treatments. For each treatment condition, traction force time courses from one representative experiment are shown for CAF spheroids in 1.2 mg/ml collagen hydrogels over 24 h. Indicated compounds or matched vehicle controls are added at measurement onset. Each line represents one spheroid, and n indicates the number of spheroids in the shown experiment.

**Extended Data Figure 14:**
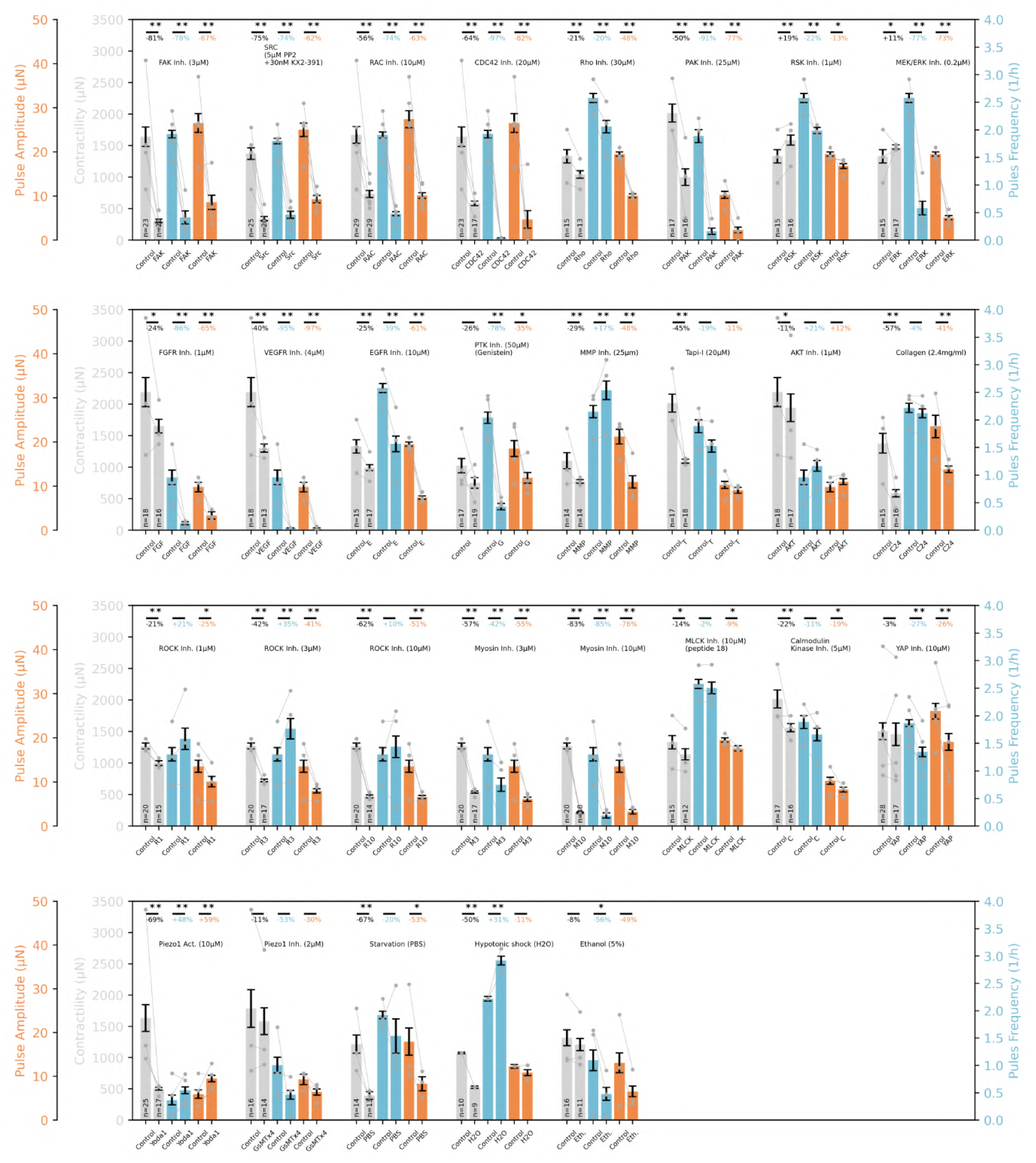
Spheroid contractility, pulse frequency, and pulse amplitude across different drug treatments. Spheroid contractility, pulse frequency, and pulse amplitude are measured in 1.2 mg/ml collagen hydrogels over 24 h. Indicated compounds or matched vehicle controls are added at measurement onset (see Methods and SI Fig.9for additional details). Bars indicate mean*±* se over all spheroids from individual measurements, and percentage values indicate the relative change between the mean values. Gray dots and lines indicate the mean of individual experiments. ** and * indicate significance of p<0.01 and p<0.05 for a student’s t-test.

**Extended Data Figure 15:**
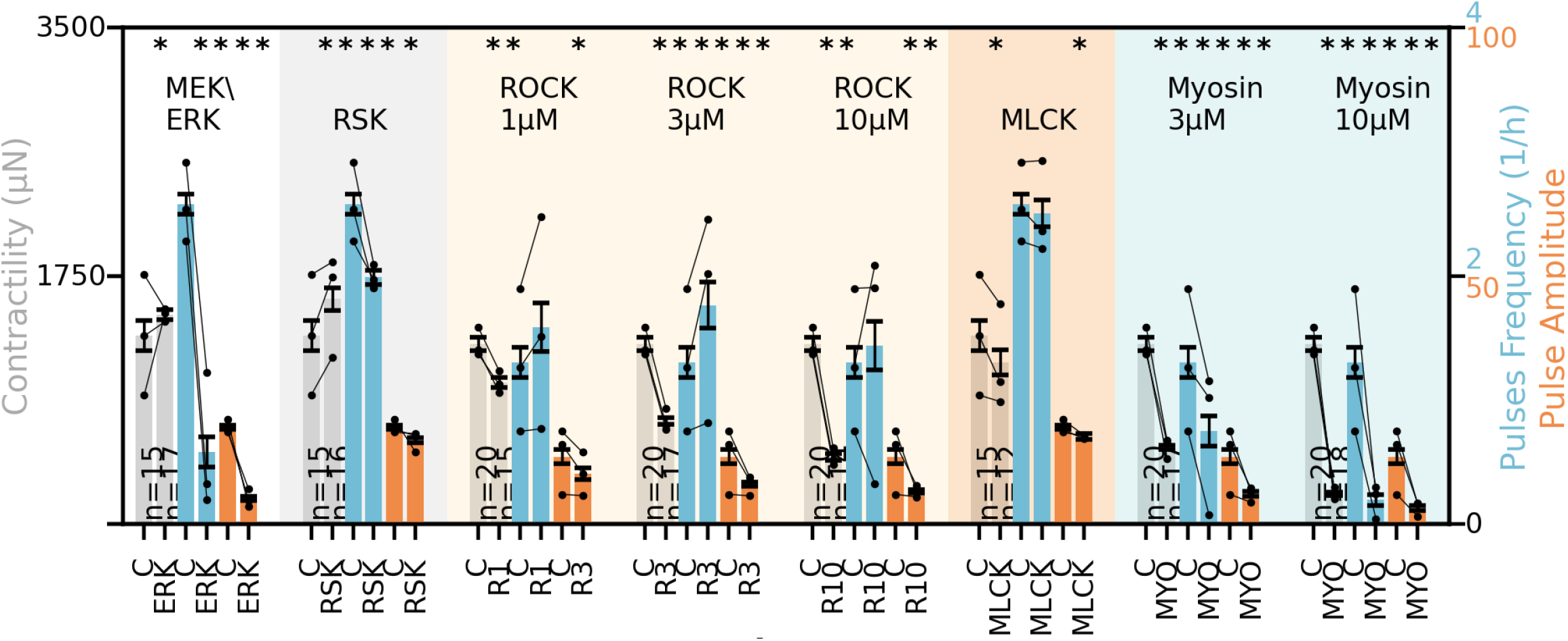
Effects of contractile machinery inhibition on spheroid contractility and force oscillations. Spheroid contractility (gray), pulse frequency (blue), and pulse amplitude (orange) are quantified over 24 hours for control (C) and drug-treated spheroids. The inhibitors target different components of the cellular contractile machinery (see Figure 4and SI Fig.9for additional details). Bars show the mean*±* se across at least three individual experiments. For each experiment, values are first averaged across all spheroids measured during the 24 h measurement period. The total number of spheroids is indicated by n, whereas black dots represent the means of individual experiments. Lines connect corresponding compound-treated and control experiments. The respective compound and molecular target are indicated above the bars. Significance is determined using a paired t-test, where * indicates p*<*0.05 and ** indicates p*<*0.01.

**Extended Data Figure 16:**
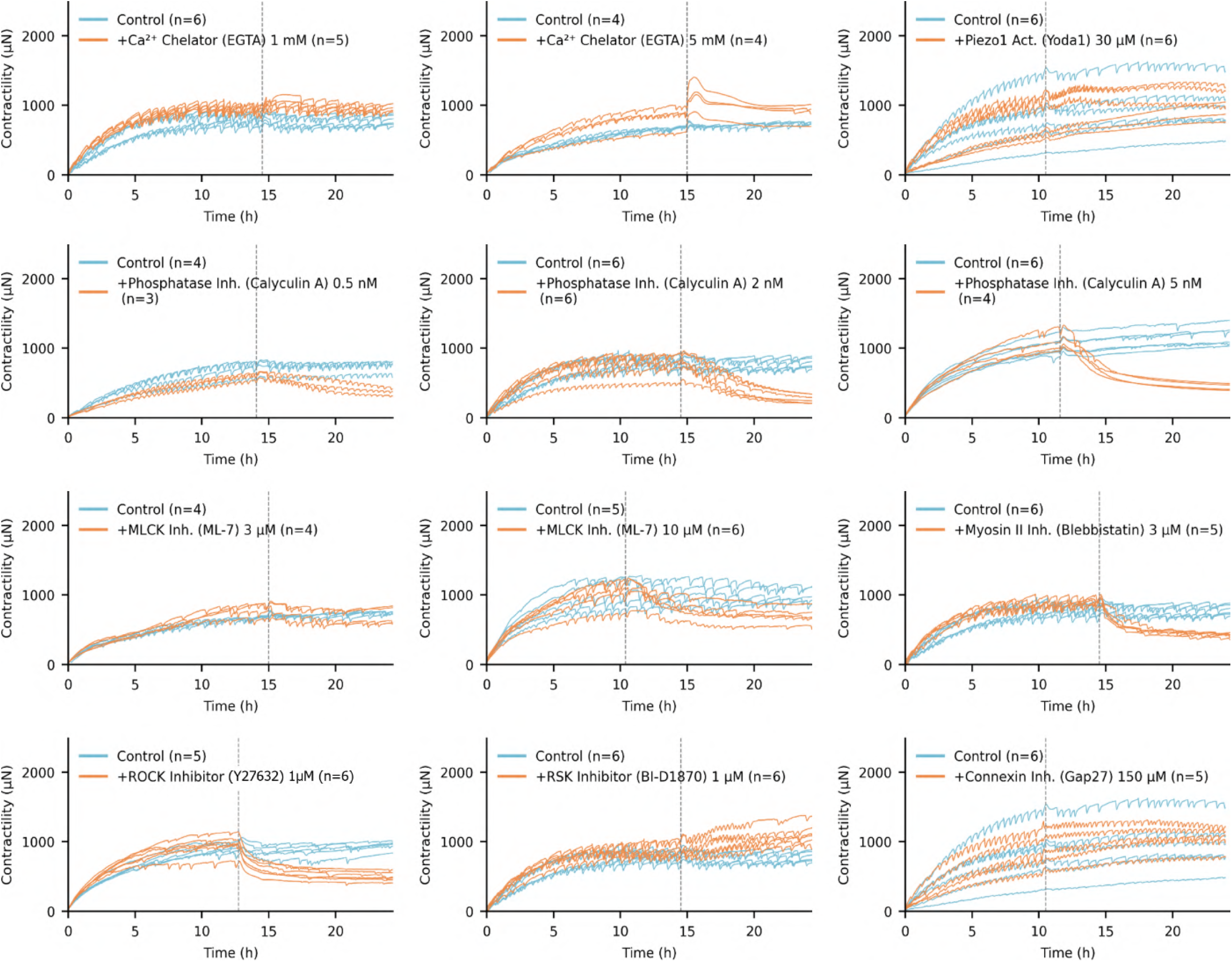
Traction force time courses for dynamic drug-screening experiments targeting calcium signaling and the contractile machinery. For each treatment condition, spheroid traction force time courses from one representative experiment are shown in 1.2 mg/ml collagen gels over 24 h. Compounds targeting calcium signaling or the contractile machinery, or the corresponding matched vehicle controls, are added between approximately 9-15 h after measurement onset, as indicated by the gray dotted lines. Each line represents an individual spheroid, and n indicates the number of spheroids in the shown experiment.

**Extended Data Figure 17:**
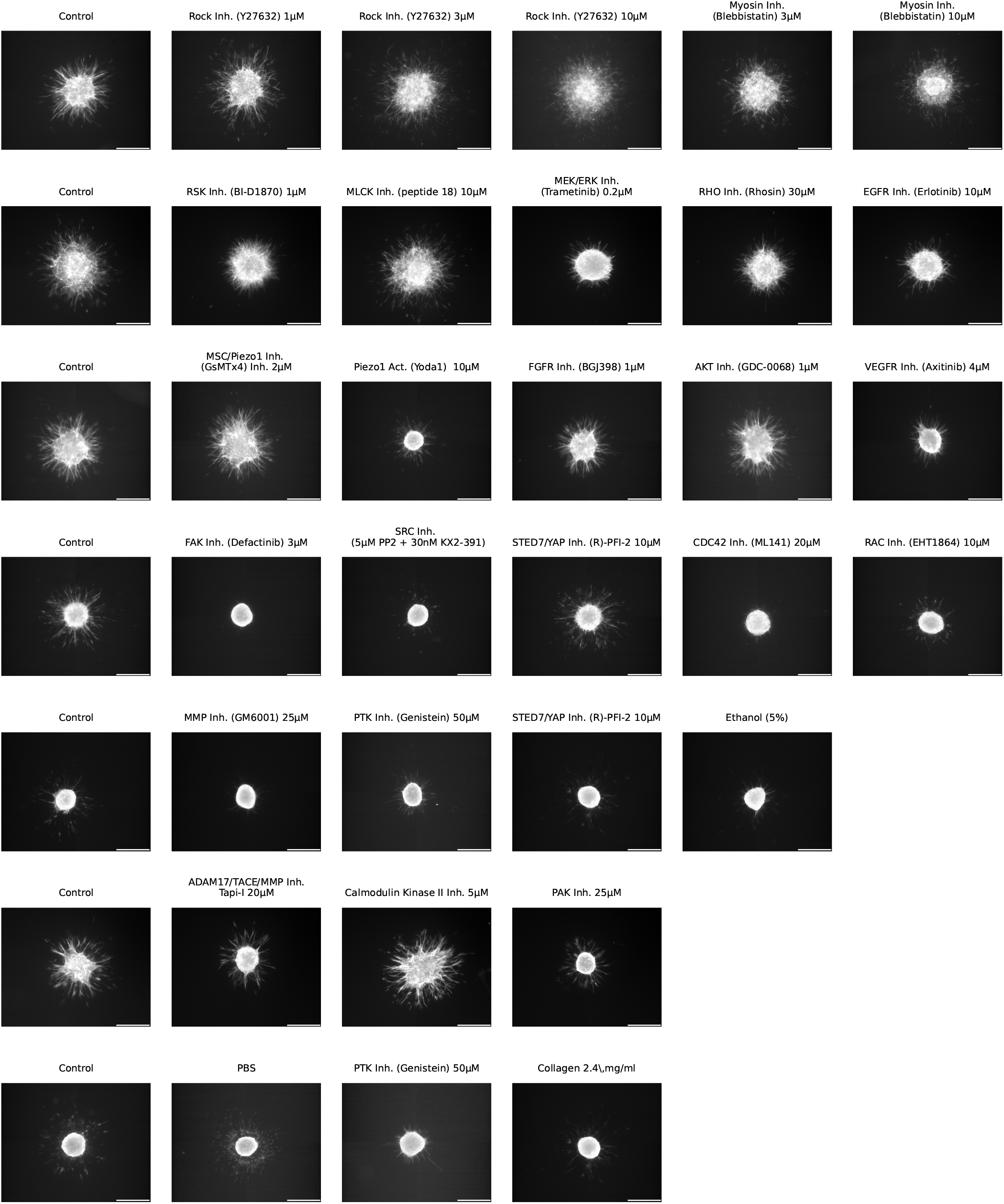
Spheroid cell invasion in collagen matrices under different drug treatments. Fluorescence images of F-actin stained with TRITC-conjugated phalloidin are acquired at the equatorial plane of representative spheroids 24 h after embedding in 1.2 mg/ml collagen hydrogels. Indicated drugs or vehicle controls are added at the onset of the 24 h incubation period (see SI Fig.9for additional details). Each image pair originates from a single experimental day, with the corresponding control spheroid shown on the left and the drug-treated spheroid on the right. The scalebar indicates 500 µm.

**Extended Data Figure 18:**
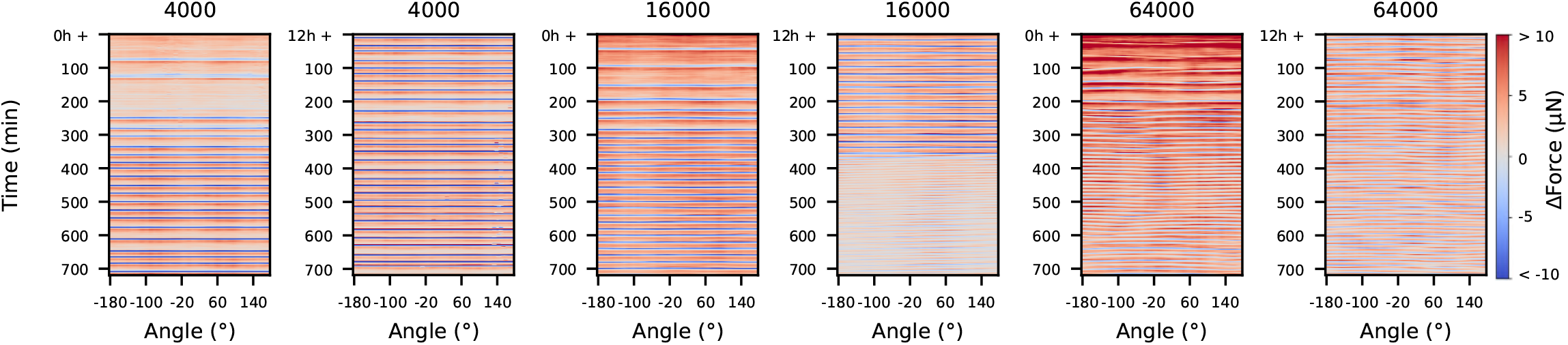
Kymographs showing spatial force changes for different spheroid sizes. The kymographs show changes in force between consecutive 1 min time points as a function of time and angular position around each spheroid. Forces are calculated in 5° angular sectors, with red and blue indicating inward- and outward-directed force changes, respectively. Spheroids are generated from 4000 cells (left), 16000 cells (middle), and 64000 cells (right), as shown inFigure 1. Force reconstruction from matrix deformations within the angular sectors is illustrated inFigure 7.

**Extended Data Figure 19:**
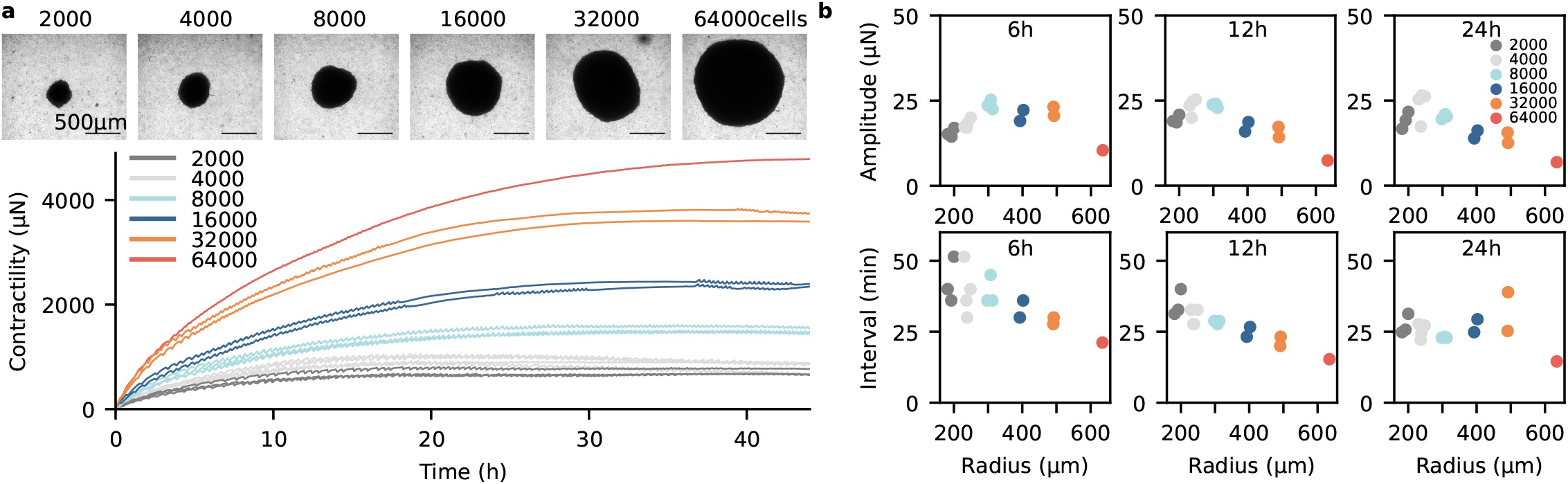
Force oscillations in different sized CAF spheroids. **a**, Brightfield microscopy images (top) and collective traction force measurements (bottom) of spheroids generated from 2000 to 64000 cells. **b**, Pulse amplitude and pulse interval of spheroids of different sizes, calculated over the first 6 hours, 12 hours, or 24 hours after embedding in collagen. Overall, spheroid contractility increases with spheroid size. As spheroid size increases, the interval between force oscillations decreases. Over time, pulse amplitudes increase in smaller spheroids but decrease in larger spheroids.

**Extended Data Figure 20:**
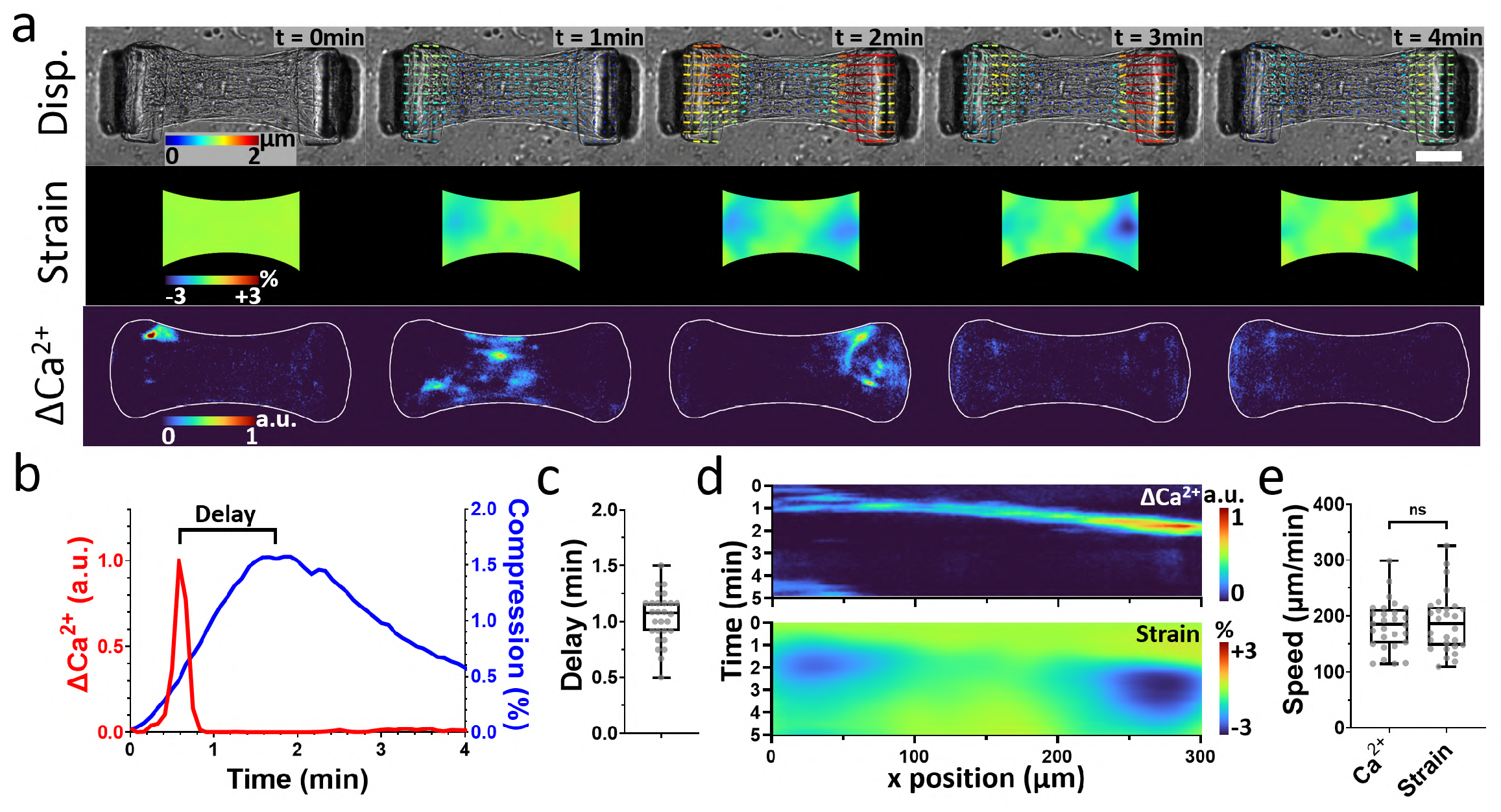
Force and calcium oscillations in microtissues. **a**, Brightfield images overlaid with matrix displacement vectors (top), strain maps (middle), and changes in the intracellular calcium signal (bottom) during force oscillations in microtissues generated from 300 patient-derived human cancer-associated fibroblasts (CAFs), isolated from a Lieberkühnian adenocarcinoma of the upper rectum. Intracellular calcium is visualized using the calcium indicator Fluo-4 AM at a concentration of 2 µM. The scale bar represents 100 µm. **b**, Time courses of the calcium signal and compressive force during a representative force oscillation. **c**, Time delay between the calcium-signal peak and the force peak, measured across 28 microtissues. **d**, Kymograph showing the propagation of calcium and strain waves over time along the horizontal axis of the tissue. **e**, Propagation speeds of calcium and strain waves measured across 28 microtissues. For each box plot, the box extends from the 25th to the 75th percentile, the central line indicates the median, the whiskers extend to the minimum and maximum values, and individual data points are overlaid.

**Extended Data Figure 21:**
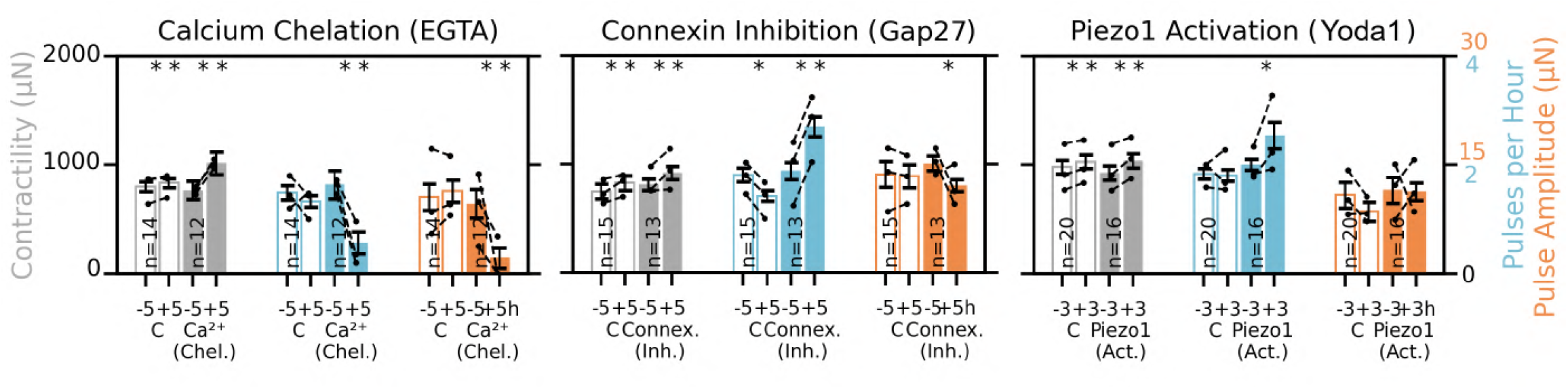
Effect of calcium inhibiting drugs on contractility and force oscillations. To investigate the role of calcium signaling, compounds that perturb calcium availability, intercellular communication, or mechanosensitive ion-channel activity are added to the medium during spheroid traction force measurements (see Figure 7 and SI Fig.9for additional details). Contractility (gray), pulse frequency (blue), and pulse amplitude (orange) are calculated for each experiment by averaging across all spheroids within a 3 h window for Yoda1 or a 5 h window for EGTA and Gap27 immediately before and after compound addition. EGTA (5 mM), Gap27 (150 µM), and Yoda1 (30 µM) are used to chelate extracellular Ca^2+^, inhibit connexin-mediated gap junction communication, and activate Piezo1, respectively. Bars show the mean *±*se across experiments, and black dots indicate the means of individual experiments. Filled bars represent experiments with compound addition, whereas unfilled bars represent the corresponding vehicle controls containing the same final solvent concentration. Compound names and their molecular targets are indicated below and above the bars, respectively. Significance is determined using a paired t-test, where * indicates p*<*0.05 and ** indicates p*<*0.01.

**Extended Data Figure 22:**
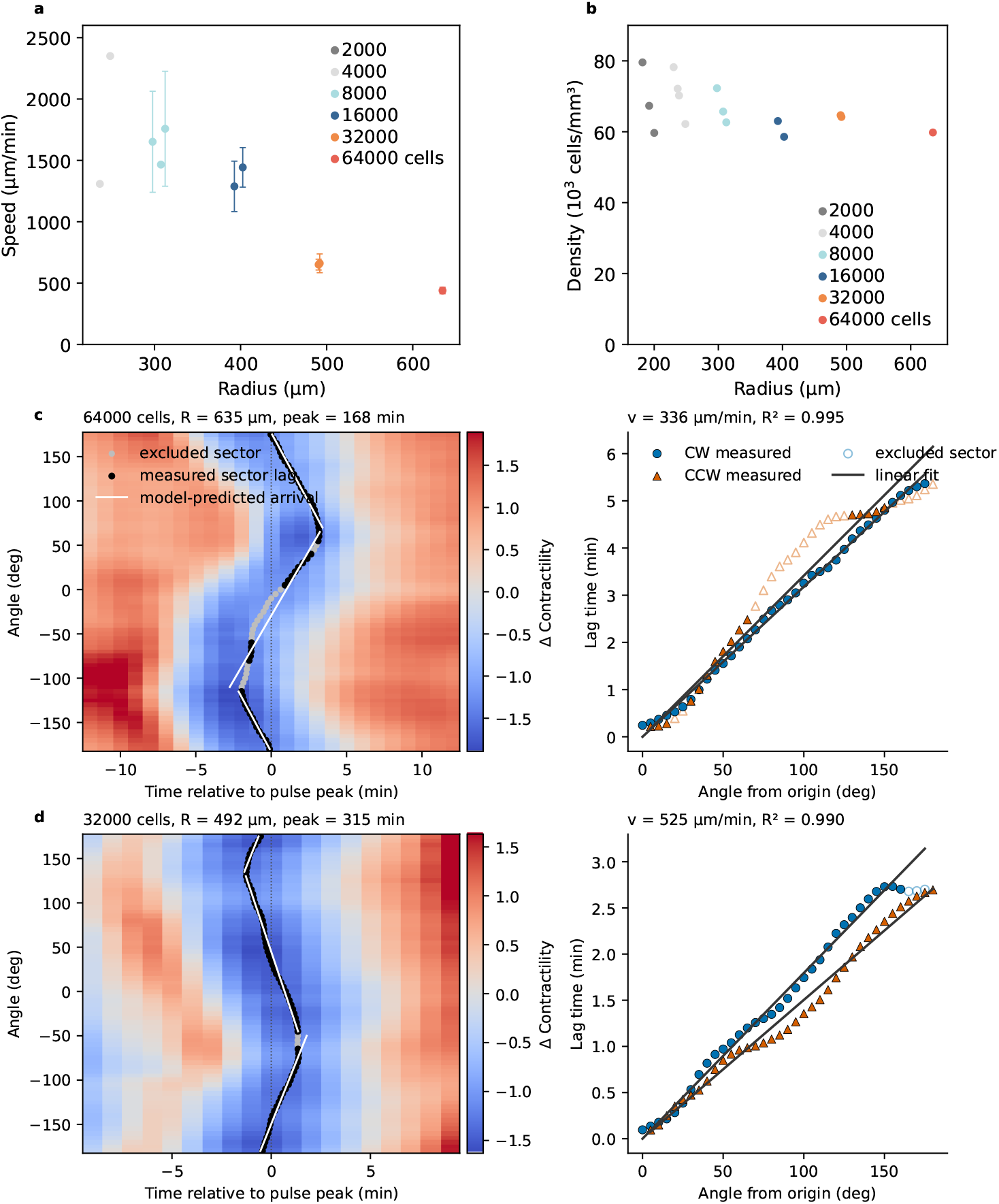
Spheroid size, cell packing density, and angle-resolved measurement of mechanical-wave speed. **a**, Mechanical-wave speed as a function of spheroid radius. Points show mean *±*se across pulses of each spheroid. Colors indicate the initial cell number. **b**, Estimated cell packing density, calculated as the initial cell number divided by the spheroid volume, assuming a spherical spheroid shape. **c**,**d**, Representative angle-resolved wave analyses for spheroids initiated with 64 000 and 32 000 cells, respectively. Left: Kymographs with predicted arrival times. Right: sector delays with clockwise (CW) and counterclockwise (CCW) fits. Contractility is measured every minute in 5° angular sectors. For each pulse, the median signal across all sectors serves as a reference curve that represents the typical pulse shape. Each angular trace is shifted in time to maximize its Pearson correlation with this reference. The required time shift defines the delay of that sector. The optimal shift is interpolated between consecutive frames, allowing delays below 1 min to be estimated. The wave origin is chosen as the angular position that gives the clearest linear relationship between delay and angular distance. CW and CCW propagation are fitted jointly with a common origin but separate slopes. In case only one wave front is visible, only one line is fitted. Strongly deviating sectors are excluded using a median-absolute-deviation criterion. Wave speed is calculated from the fitted slopes (delay per angle) and the spheroid radius. Pulses are included when the median of the maximum Pearson correlation coefficients between the sector traces and the reference is at least 0.90, the linear relationship between sector delay and angular distance has an *R*^2^ of at least 0.90, the CW and CCW fits show consistent propagation and their speeds differ by no more than 2.5-fold, and the fitted delay over 150° is at least 0.25 min. In the kymographs, black points indicate included sectors, gray points indicate excluded sectors, and white lines show the predicted arrival times. In the delay plots, filled blue circles show included CW sectors and filled orange triangles show included CCW sectors. Open symbols indicate excluded sectors, and solid black lines show the linear fits. Numbers of analyzed pulses per spheroid are: 4 000 cells, 1/1; 8 000 cells, 1/4/4; 16 000 cells, 9/5; 32 000 cells, 9/10; and 64 000 cells, 12 (56 pulses in total. No pulses met the inclusion criteria for 2 000-cell spheroids).

**Extended Data Figure 23:**
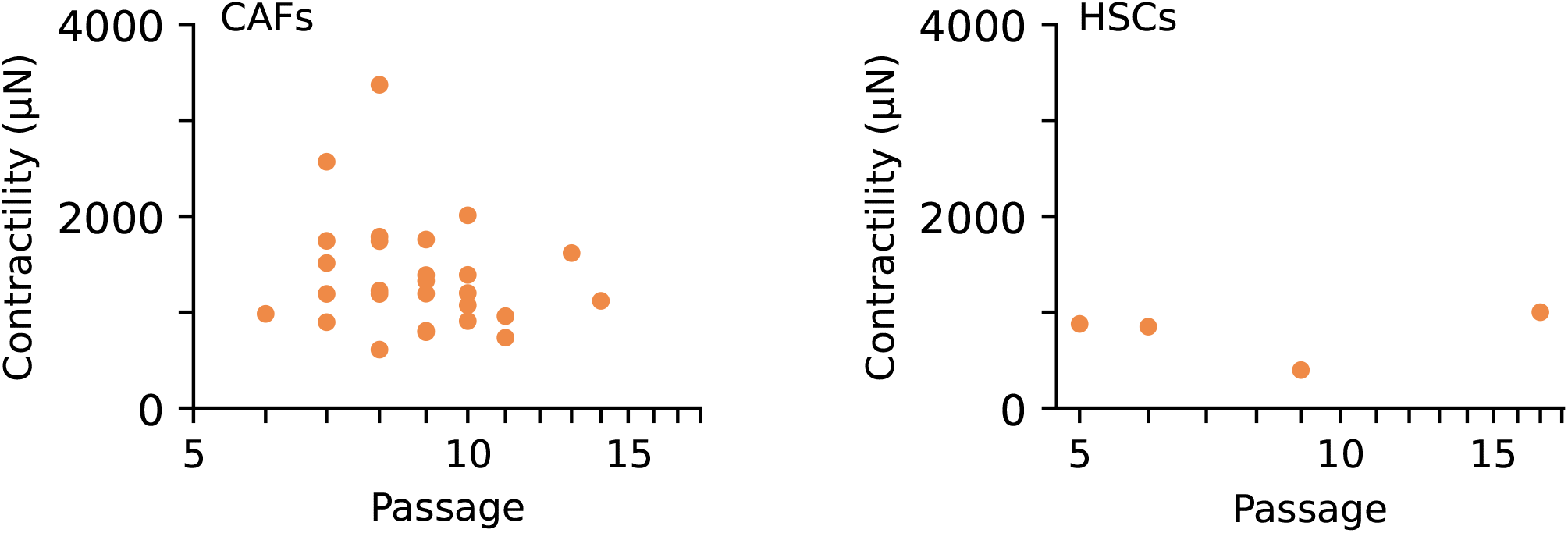
Spheroid contractility as a function of cell passage number. Mean contractility of individual spheroids measured over 24 h in collagen (typically 8–12 spheroids per experiment) as a function of the cell passage number used for the corresponding traction force measurement. Data for breast cancer-associated fibroblasts are shown on the left, whereas data for hepatic stellate cells are shown on the right. No systematic dependence of spheroid contractility on cell passage number is observed.

**Extended Data Figure 24:**
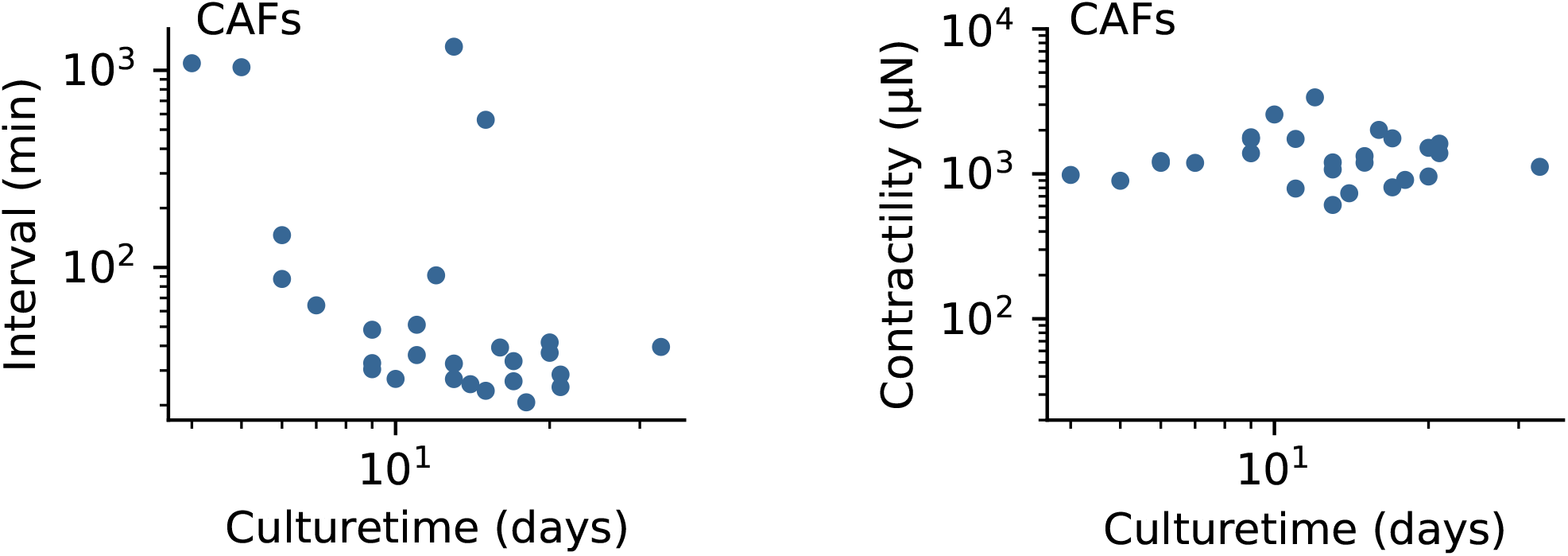
Spheroid contractility and pulse interval as a function of cell culture time. Mean spheroid contractility and time interval between force oscillations in breast cancer-associated fibroblast (CAF) spheroids, measured over 24 h in collagen (typically 8–12 spheroids per experiment), as a function of the time for which the cells are maintained in culture after thawing and before spheroid generation. The interval between force oscillations decreases with increasing time in culture after thawing, corresponding to a higher pulse frequency, whereas spheroid contractility shows no systematic dependence on time in culture after thawing.

**Extended Data Figure 25:**
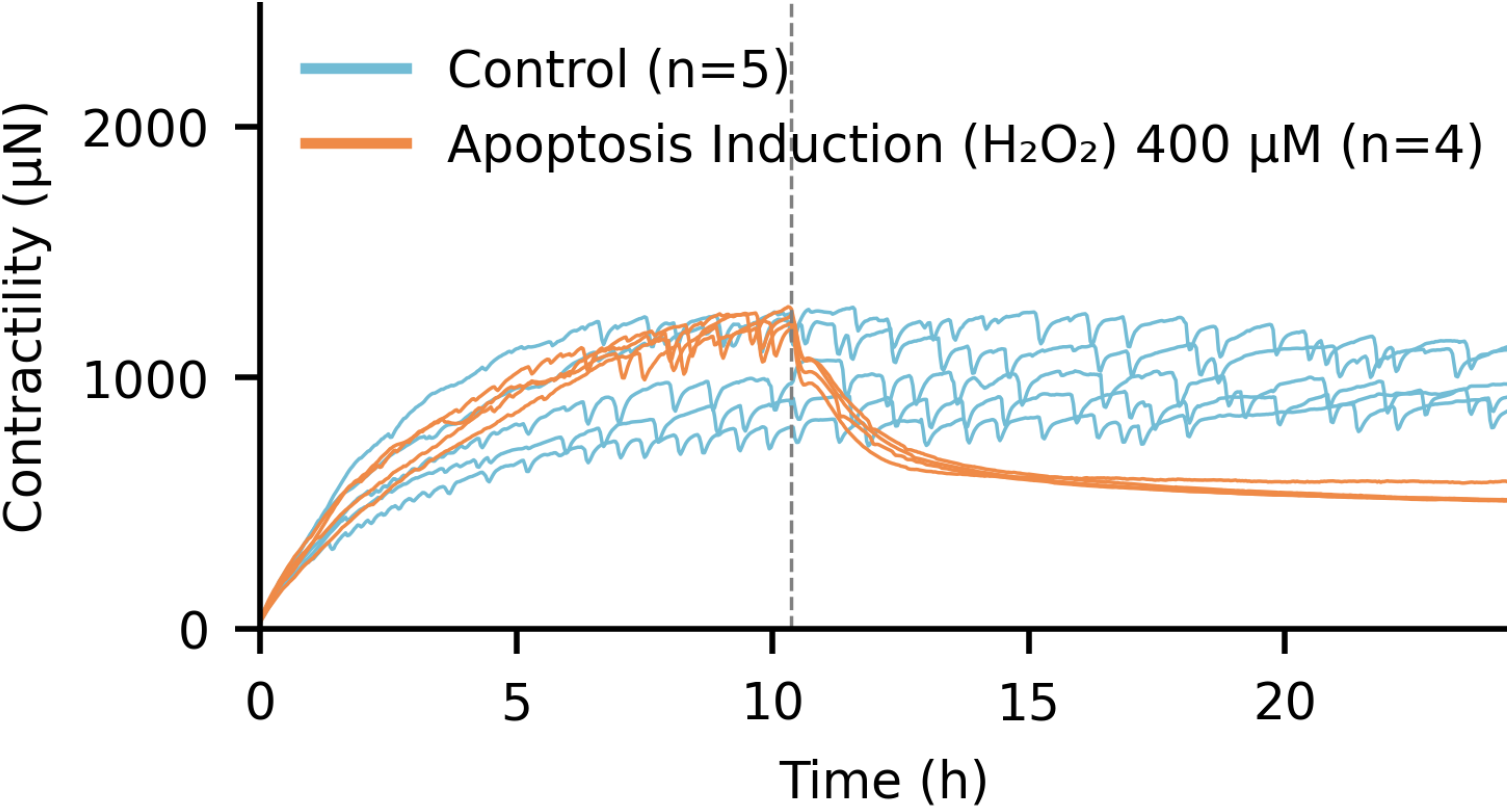
Effect of hydrogen peroxide treatment on spheroid contractility. Spheroid traction force time courses are measured in 1.2 mg/ml collagen gels over 24 h for control spheroids and spheroids treated with 400 µM hydrogen peroxide (H_2_O_2_) to induce oxidative stress and apoptosis. The gray dotted line indicates the time of H_2_O_2_ addition, and n indicates the number of individual spheroids shown.

**Supplementary Video 1: Force oscillations in a CAF spheroid in 1.2 mg/ml collagen over 24 h**. Video showing a spheroid generated from 4000 primary human Luminal B CAFs and embedded in a 3D collagen matrix (1.2 mg/ml). Brightfield images are acquired at the equatorial plane of the spheroid every 2 min over 24 h. Microbeads (4 µm) are used to trace cell-generated matrix deformations within the collagen.

**Supplementary Video 2: Force oscillations in a healthy breast fibroblast spheroid in 1.2 mg/ml collagen over 24 h**.

Video showing a spheroid generated from 4000 primary human fibroblasts isolated from healthy breast tissue and embedded in a 3D collagen matrix (1.2 mg/ml). Brightfield images are acquired at the equatorial plane of the spheroid every 2 min over 24 h. Microbeads (4 µm) are used to trace cell-generated matrix deformations within the collagen.

**Supplementary Video 3: Cumulative matrix deformations around CAF spheroid in 1.2 mg/ml collagen**.

Video showing cell-generated cumulative matrix deformations within the equatorial plane of a luminal B spheroid in 3D collagen over a 24 hour period. Matrix deformations are tracked from timelapse images using 4 µm-sized microbeads and particle image velocimetry (26,32).

**Supplementary Video 4: Time-resolved matrix deformations around a CAF spheroid in 1.2 mg/ml collagen**

The video shows the direction of consecutive matrix deformations within the equatorial plane of a breast cancer-associated fibroblast spheroid derived from a luminal B breast carcinoma during 24 h in 3D collagen. Force oscillations are visible as brief periods of outward-directed matrix deformation. Matrix deformations are quantified from the timelapse images by particle image velocimetry using embedded 4 µm-sized microbeads (26,32).

**Supplementary Video 5: Force oscillations in an HSC spheroid embedded in 1.2 mg/ml collagen over 24 h**.

The video shows a spheroid generated from 4000 hepatic stellate cells (HSCs) embedded in a 3D collagen matrix (1.2 mg/ml). Brightfield images are acquired at the spheroid equatorial plane every 5 minutes over a period of 24 h. Embedded 4 µm-sized microbeads are used to track cell-generated matrix deformations within the collagen. Force oscillations emerge with intervals of approximately 20-30 minutes.

**Supplementary Video 6: Force oscillations in a microtissue generated from 4000 CAFs over 25 h**.

The video shows 4000 breast cancer-associated fibroblasts (CAFs) derived from a luminal B breast carcinoma and embedded in 0.6 mg/ml collagen self-assembling into a microtissue between two PDMS pillars, as described previously (28). Brightfield images are acquired in the focal plane at the top of the pillar array every 5 minutes over a period of 25 h. After self-assembly, the cells progressively generate contractile forces that deflect the two pillars towards each other. Spontaneous oscillatory pillar deflections become visible after approximately 15 hours.

**Supplementary Video 7: Collagen fiber dynamics during force oscillations**.

The video shows two neighboring spheroids, each generated from 4000 breast cancer-associated fibroblasts (CAFs) derived from a luminal B breast carcinoma, embedded in a 3D collagen matrix (1.2 mg/ml). Confocal reflection images of the collagen fiber network in the equatorial plane of the spheroids are acquired every 1 minute using an upright laser-scanning confocal microscope (Leica SP5, Leica, Wetzlar). Over the course of the measurement, the spheroids gradually move towards the bottom of the dish. No visible rupture or discontinuity of the collagen fiber network is observed during the force pulses.

**Supplementary Video 8: Local collagen stiffening around a contractile spheroid**.

The video shows the time-dependent local stiffness of the collagen matrix around a spheroid generated from 4000 breast cancer-associated fibroblasts (CAFs) derived from a luminal B breast carcinoma. The local strain-dependent collagen stiffness is inferred from the measured matrix deformations using our nonlinear elastic material model of collagen fiber networks (33,34), parameterized using shear-rheological and extensional-strain measurements (34).

**Supplementary Video 9: Force oscillations in different matrices**.

The video shows spheroids generated from 4000 breast cancer-associated fibroblasts (CAFs) derived from a luminal B breast carcinoma, embedded in different matrices and imaged over 24 h or 48 h. Embedded 4 µm-sized microbeads are used to track cell-generated matrix deformations. The hydrogels include collagen at different concentrations, two GelMA formulations with different stiffnesses and approximately linear elasticity, and Matrigel, which exhibits weaker strain stiffening than collagen. See SI Fig.6 for the rheological characterization of the different materials. Matrigel samples display fluid-filled inclusions at the interface between the two hydrogel layers, which are not observed in the other materials. The scale bar represents 500 µm. Collagen gels, which exhibit nonlinear strain stiffening, show strong spheroid contractility and pronounced force oscillations.

**Supplementary Video 10: Comparison of force oscillations in collagen and alginate-based hydrogels**.

The video shows spheroids generated from 4000 breast cancer-associated fibroblasts (CAFs) derived from a luminal B breast carcinoma, embedded in different matrices and imaged over 24 h or 48 h. Embedded 4 µm-sized microbeads are used to track cell-generated matrix deformations. The matrices are collagen hydrogels (1.2 mg/ml) and alginate dialdehyde-gelatin (ADA-GEL) hydrogels containing 2.5% ADA and 2.5% GEL, as described previously (62,63). Images of the ADA-GEL samples are stabilized to correct for swelling-induced bulk drift. The scalebar represents 300 µm. The nonlinear strain-stiffening collagen gels support strong spheroid contractility and pronounced force oscillations, whereas contractility and oscillatory matrix deformations are reduced in ADA-GEL.

**Supplementary Video 11: Effects of myosin II and ROCK inhibition on spheroid invasion in collagen**.

The video shows spheroids generated from 4000 breast cancer-associated fibroblasts (CAFs) derived from a luminal B breast carcinoma, embedded in 1.2 mg/ml collagen matrices and imaged over 24 h. Embedded 4 µm-sized microbeads are used to track spheroid-generated matrix deformations. Spheroids treated with the myosin II inhibitor blebbistatin (10 µM) or the Rho-associated kinase (ROCK) inhibitor Y-27632 (10 µM) show predominantly sheet-like invasion. In contrast, the more contractile control spheroids show strand-like invasion along aligned collagen fibers (SI Fig.2).

**Supplementary Video 12: Effects of MEK/ERK inhibition on CAF spheroids in collagen**

The video shows spheroids generated from 4000 breast cancer-associated fibroblasts (CAFs) derived from a luminal B breast carcinoma, embedded in 1.2 mg/ml collagen matrices and imaged over 24 h. Embedded 4 µm-sized microbeads are used to trace spheroid-generated matrix deformations. Spheroids are treated with trametinib (0.2 µM) or the corresponding vehicle control. Inhibition of MEK/ERK signaling strongly reduces spheroid force oscillations.

**Supplementary Video 13: Spheroid force oscillations synchronize through a 3D collagen matrix**

**Top**, Brightfield microscopy videos of a closely spaced spheroid pair (left) and a more separated spheroid pair (right). Each spheroid is generated from 4000 breast cancer-associated fibroblasts (CAFs) derived from a luminal B breast carcinoma and embedded in a 3D collagen matrix at a concentration of 1.2 mg/ml. **Bottom**, Corresponding high-pass-filtered contractile-pressure time courses for the closely spaced pair (left) and the more distant pair (right), showing that the force oscillations within each spheroid pair synchronize over time.

**Supplementary Video 14: Force oscillations within CAF spheroids of different sizes**

Brightfield timelapse videos show force oscillations in breast cancer-associated fibroblast (CAF) spheroids generated from 4000, 16000, 32000, or 64000 cells and embedded in 3D collagen gels over 20 h.

**Supplementary Video 15: Force oscillations and angle-resolved contractile pressure in CAF spheroids of different sizes**

**Top**, Brightfield timelapse videos show force oscillations in breast cancer-associated fibroblast (CAF) spheroids generated from 2000, 4000, 8000, 16000, 32000, or 64000 cells and embedded in 3D collagen gels over 10 h. **Bottom**, Corresponding angle-resolved contractile pressure around each spheroid. Matrix displacement fields are quantified from the brightfield images using particle image velocimetry (26,32). The contractile pressure associated with each matrix displacement is reconstructed from its amplitude and radial distance from the spheroid center, normalized to spheroid size (26). For each time point, the pressure estimates are averaged within 5° angular sectors around the spheroid, showing the spatial propagation of mechanical waves during force pulses.

**Supplementary Video 16: Traveling mechanical wave during a force oscillation in a 32000-cell CAF spheroid**.

The video shows force oscillations in a breast cancer-associated fibroblast (CAF) spheroid generated from 32000 cells and embedded in a 3D collagen gel. Matrix displacement fields between consecutive images are quantified using particle image velocimetry. The colormap and arrows indicate the direction of matrix displacement, with red representing inward-directed and blue representing outward-directed displacements. During individual force oscillations, local matrix deformations propagate around the spheroids as a mechanical wave.

**Supplementary Video 17: Mechanical wave propagation during force oscillations in CAF spheroids of different sizes**.

Brightfield timelapse videos show force oscillations in breast cancer-associated fibroblast (CAF) spheroids generated from 16000, 32000, or 64000 cells and embedded in 3D collagen gels over 10 h. Matrix displacement fields between consecutive images are quantified using particle image velocimetry. The colormap and arrows indicate the direction of matrix displacement, with red representing inward-directed and blue representing outward-directed displacements. During individual force oscillations, local matrix deformations propagate around the spheroids as a mechanical wave.

**Supplementary Video 18: Transition from pulsatile force oscillations to a traveling mechanical wave in a 16000-cell CAF spheroid**.

The video shows force oscillations in a breast cancer-associated fibroblast (CAF) spheroid generated from 16000 cells and embedded in a 3D collagen gel. Matrix displacement fields between consecutive images are quantified using particle image velocimetry. The colormap and arrows indicate the direction of matrix displacement, with red representing inward-directed and blue representing outward-directed displacements. During individual force oscillations, local matrix deformations propagate around the spheroids as a mechanical wave. After approximately 18 h, the spatial pattern of the pulsatile force oscillations transitions into a traveling mechanical wave that propagates around the spheroid periphery. See also the force kymographs in SI Fig.18.

**Supplementary Video 19: Calcium waves propagating within a CAF spheroid**

Timelapse imaging during a force oscillation in a spheroid generated from human breast cancer-associated fibroblasts (CAFs), acquired at the spheroid equatorial plane. **Left**, Confocal reflection images showing the spheroid and the surrounding 3D collagen matrix. **Right**, Intracellular calcium signal measured using 4 µM Fluo-8 AM. During the force oscillation, the calcium signal propagates across the spheroid as a wave.

**Supplementary Video 20: Calcium waves propagating within a CAF microtissue**

Timelapse imaging during a force oscillation in a microtissue generated from human cancer-associated fibroblasts isolated from a rectal adenocarcinoma. Brightfield images (**top**), intracellular calcium signals measured using 2 µM Fluo-4 (**middle**), and frame-to-frame changes in the calcium signal calculated between consecutive images (**bottom**) are shown. The calcium signal shows wave-like propagation across the microtissue during the force oscillation.

